# Label-retention expansion microscopy

**DOI:** 10.1101/687954

**Authors:** Xiaoyu Shi, Qi Li, Zhipeng Dai, Arthur A. Tran, Siyu Feng, Alejandro D. Ramirez, Zixi Lin, Xiaomeng Wang, Tracy T. Chow, Jiapei Chen, Dhivya Kumar, Andrew R. McColloch, Jeremy F. Reiter, Eric J. Huang, Ian B. Seiple, Bo Huang

## Abstract

Expansion microscopy (ExM) increases the effective resolving power of any microscope by expanding the sample with swellable hydrogel. Since its invention, ExM has been successfully applied to a wide range of cell, tissue and animal samples. Still, fluorescence signal loss during polymerization and digestion limits molecular-scale imaging using ExM. Here we report the development of label-retention ExM (LR-ExM) with a set of trifunctional anchors that not only prevent signal loss but also enable high-efficiency labeling using SNAP and CLIP tags. We have demonstrated multicolor LR-ExM for a variety of subcellular structures. Combining LR-ExM with super-resolution Stochastic Optical Reconstruction Microscopy (STORM), we have achieved molecular resolution in the visualization of polyhedral lattice of clathrin-coated pits *in situ*.

## INTRODUCTION

By physically expanding the sample before image acquisition, ExM has enabled the use of a conventional confocal microscope to achieve ∼ 70 nm lateral spatial resolution(*1-4*). Recent efforts have further enhanced the resolution of ExM either by increasing the volume expansion ratio (*5-7*) or by combining ExM with super-resolution microscopy such as Structured Illumination Microscopy (SIM) (*8-10*) STimulated Emission Depletion (STED) microscopy (*11-14*). In all these cases, the homogenization of the specimen through either protease digestion (*1-3*) or protein denaturation (*3, 4*) is essential to achieve isotropic expansion without structural distortion. To retain the spatial information of the target structures, the biomolecules of interest (e.g. protein(*2-4*), RNA (*15*)) and/or labels (e.g. dye-labeled DNA(*1*), dye-labeled antibodies (*2, 3*) or fluorescent proteins (*3, 4*)) are anchored to the hydrogel matrix prior to digestion or denaturation. Nevertheless, digestion and denaturation cause incompletely anchored proteins or protein fragments to be washed out (*2, 16, 17*), the polymerization to make the hydrogel produces free radicals that readily destroy fluorophores(*1, 3, 18*), and both factors can damage fluorescent proteins. Consequently, more than 50% of the target molecules can lose labeling after expansion(*3, 18*), which is a major issue of current ExM methods. The label loss problem is exacerbated when aiming for higher spatial resolution. First, ensuring isotropic expansion at nanometer scale requires more thorough proteinase digestion or denaturation, hence more labeled antibodies are washed out (*1-3*). Second, super-resolution microscopy often requires certain dyes that do not survive hydrogel polymerization. For example, Alexa Fluor (AF) 647, one of the best fluorophores for STORM and PALM, suffers > 90% loss of the fluorescent molecules after polymerization and digestion(*2, 3, 14*).

When the spatial resolution approaches the molecular scale, high-efficiency labeling of target molecules becomes a distinctly new requirement, as previously recognized in the development and application of super-resolution microscopy(*19-21*). This requirement is because information from unlabeled target molecules is permanently lost. Although amplifying retained labels by amplifying DNA-barcoded antibody such as isHCR(*22*) and Immuno-SABER (*23*), or using biotin (*2, 14*)) can significantly enhance the brightness, it cannot recover the lost positional information from washed-out antibodies. These methods result in bright but still incompletely labeled structures. Moreover, the bulky labels will introduce localization displacement that is detectable with the molecular resolution. On the other hand, post-expansion immunostaining can avoid dye loss during the gelation and digestion steps, such as Magnified Analysis of Proteome (MAP)(*4*), CUBIC-X (*24*) and SHIELD(*25*). Nevertheless, post-gel immunostaining can be problematic for certain targets such as neuroligin and GAP, because these proteins are damaged by denaturation and cannot be recognized by antibodies after expansion (*4*). The damage and loss of target proteins will cause underlabeling. In fact, the density of labeled targets fundamentally sets the lower limit of effective spatial resolution (*19, 26, 27*).

With the combination of ExM and super-resolution microscopy, fluorophores can theoretically be localized with the target proteins with molecular resolution. However, in practice, most ExM methods uses indirect immunostaining to label target proteins. Although convenient, immunostaining ExM has two practical problems. First, high quality antibodies are not always available. Second, bulky primary and secondary antibodies can add up to 20 nm distance between the target protein and the dye molecule or barcoding DNA strand (*5*), which is larger than the resolution of ExSTED and ExSTORM. Therefore, immunostaining is insufficient to reveal biological structures with molecular resolution. An effective solution is to express enzymatic protein tags, such as SNAP and CLIP tags, which can be recognized by highly specific and efficient fluorescent ligands (*19, 28, 29*). These two tags are about 2 nm long, much smaller than antibodies and even smaller than fluorescent proteins.

Herein we report Label-Retention Expansion Microscopy (LR-ExM), a method that solves the signal loss problem with a set of small molecule trifunctional anchors that are inert to polymerization, digestion and denaturation and that can be fluorescently labeled after expansion. We have developed this method not only for immunofluorescence labeling but also for SNAP and CLIP tags that are particularly advantageous in their small size and high labeling efficiency by organic dyes with tag-recognizing ligands.

## RESULTS

The workflow of LR-ExM includes five steps, labeling target molecules with trifunctional anchors, forming *in situ* hydrogel with cells or tissues, proteinase digestion, post-expansion fluorescence staining, and expansion (Fig. 1a). Specifically, we designed and synthesized trifunctional anchors with three arms (Fig. 1b): (1) N-hydroxysuccinimide (NHS) for connection to antibodies, benzylguanine (BG) for SNAP-tag recognition, or benzylcytosine (BC) for CLIP-tag recognition; (2) methacrylamide (MA) group for anchoring to the polymer matrix; (3) biotin or digoxigenin (DIG) as two orthogonal reporter handles for conjugation to an organic dye after expansion. We have chosen these functional groups and the molecular scaffold so that the anchors are resistant to both major fluorescence-loss causes, proteinase digestion and acrylamide gel polymerization. Therefore, target molecules labeled with the trifunctional anchors retain high labeling efficiency. The two orthogonal reporter handles enable two-color labeling and imaging. The structures and syntheses of the anchors are described in detail in Fig. S1 and S2 and Supplementary Materials and Methods.

**Fig. 1.**
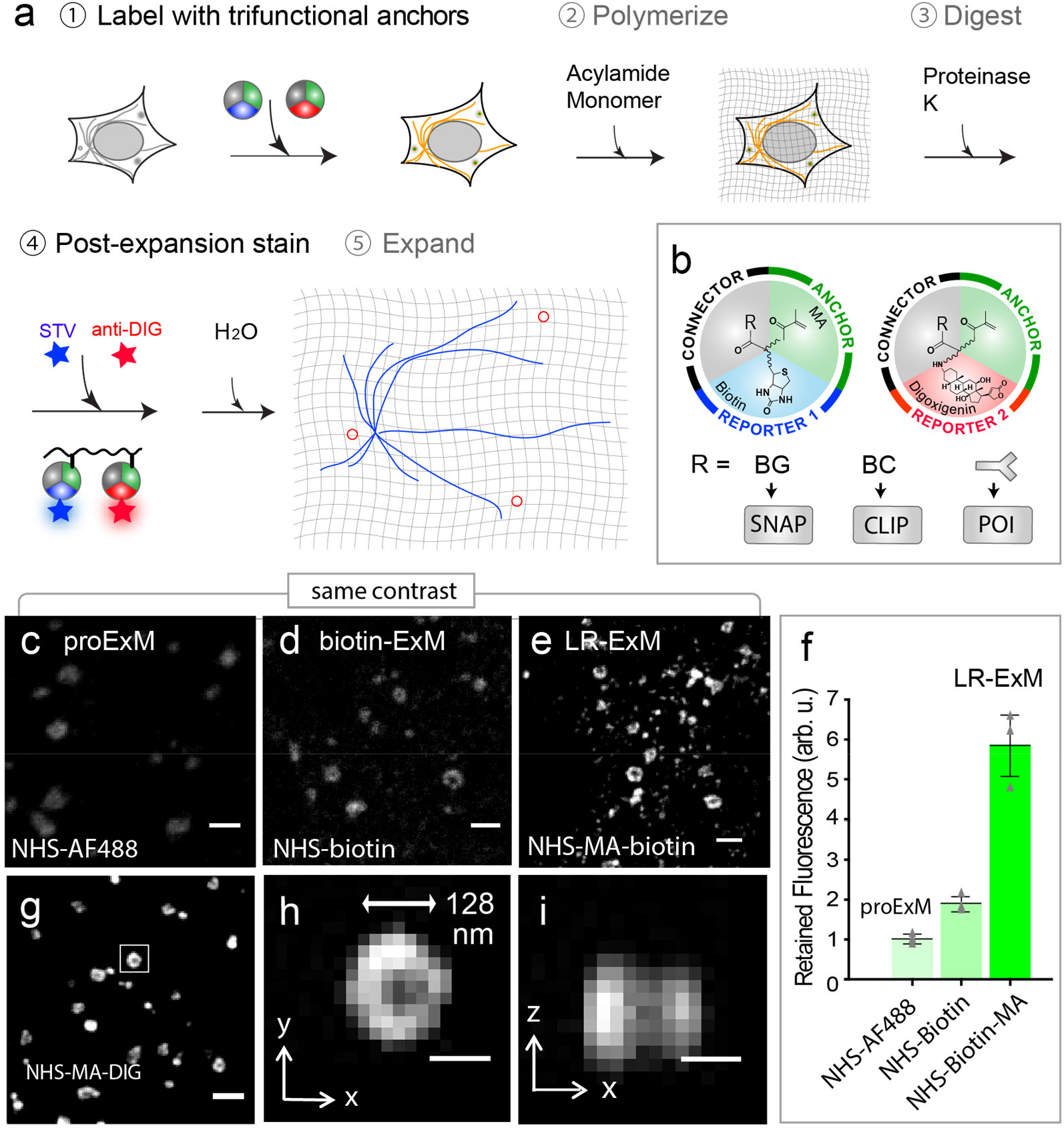
Workflow and characterization of LR-ExM. (a) Workflow of LR-ExM. (b) Schematic of trifunctional anchors. (c-e) ExM confocal images of CCPs in U2OS cells indirectly immunostained for α tubulin. (c) proExM using AF488-conjugated secondary antibodies. (d) ExM with post-expansion labeling using Biotin-conjugated antibodies. (e) LR-ExM using antibodies conjugated with NHS-MA-biotin tri-functional anchor. Samples for (d) and (e) were post-expansion stained with streptavidin-AF488. (f) Intensity quantification of (c-e). Error bars are standard deviations. N = 3 for each case. (g) LR-ExM confocal image of CCPs in U2OS cells immunostained indirectly with secondary antibodies conjugated with NHS-MA-DIG anchor, post-expansion stained with anti-Digoxigenin antibody. (h, i) Cross sections of the CCP in the boxed area of (g). The length expansion ratios for images (c), (d), (e), and (g/h/i) are 4.3, 4.5, 4.6, and 4.3, respectively. The length expansion ratio for samples used in plot (f) is 4.5 ± 0.2. Scale bars, 500 nm for (c-e, g), and 100 nm for (h, i). All scale bars in pre-expansion units.

With different trifunctional anchors, LR-ExM is compatible with both immunofluorescent and enzymatic protein tags (Fig. 1a). For immunofluorescence, we stained fixed cells with antibodies conjugated to NHS-MA-biotin or NHS-MA-DIG, proceeded with the standard ExM procedure of gel polymerization and proteinase K digestion, then stained the gel with fluorescently labeled streptavidin for biotin anchor and/or anti-DIG antibody for before expanding the gel in water and imaging. For labeling with SNAP or CLIP tag, the procedure is nearly identical except that fixed cells were directly treated with BG- or BC-trifunctional anchors.

To demonstrate label retention, we compared ExM of U2OS cells immunostained for microtubules and clathrin heavy chain using secondary antibodies conjugated to Alexa Fluor 488 (AF488) dye (following the proExM protocol (*3*)), biotin (proExM followed by post-expansion staining with AF488-streptavidin (*2, 14*)) or NHS-MA-biotin (our LR-ExM). Fig. 1c-e showcased the clathrin-coated pit (CCP) images (Fig. 1c-e) processed with these three kinds of secondary antibodies, with the same contrast. We conjugated streptavidin with an average dye: protein ratio of 1:1 so that the fluorescence level in the three cases can be directly compared. On average the LR-ExM generated a fluorescence signal that is 5.8 ± 0.8 (mean ± standard deviation, N = 3) times as high as that by AF488 antibody (Fig. 1f). For quantification methods, see Supplementary Information and Fig. S3). The biotin-antibody sample generated 1.9 ± 0.2 (mean ± standard deviation, N = 3) times of the fluorescence signal compared to proExM. Taking these values together, we concluded that out of the ∼ 83% label loss by proExM, ∼ 15% can be attributed to polymerization and ∼ 68% to digestion. These numbers may be rescaled since the biotin may not be completely conserved. The fraction of fluorescence loss caused by digestion is dependent on the labeling tags such as antibody, streptavidin, and GFP, and is affected the digestion condition such as digestion duration, buffer, and temperature. Differences between the protocols may lead to contradictory results. In this evaluation, we focused on IgG antibodies, and followed digestion condition of proExM(*3*). See methods section and Fig. S3 for detailed procedure.

We calibrated the length expansion ratio of our LR-ExM protocol to range from 4.3 to 4.7. Fitting the cross-section profiles of microtubules to a Gaussian peak then gave a full width at half maximum (FWHM) of 84 ± 1.3 nm (mean ± standard deviation, N = 3 independent experiments) after rescaling by the expansion ratio. Taking the size of immunostained microtubules (*30*) into the cross-section profile for fitting (*1, 31*), we obtained an effective resolution of 71 ± 1.6 nm. At this effective resolution, we could resolve the hollow shape of CCPs (Fig. 1h and i). Using anti-DIG antibody with a high dye:protein ratio (10:1) produces brighter signal (Fig. 1g), though we do not expect obvious differences in the actual labeling efficiency.

We demonstrated two-color LR-ExM by co-immnunostaining CCPs and microtubules with antibodies conjugated with NHS-MA-DIG and NHS-MA-biotin, respectively (Fig. 2a, b). Similarly, we demonstrated LR-ExM for SNAP-tag and CLIP-tag with BG-MA-biotin and BC-MA-DIG trifunctional anchors, respectively (Fig. 2c for CCPs by overexpressing SNAP-clathrin and Fig. 2d for mitochondria by TOMM20-CLIP), including two-color imaging using both enzymatic protein tags owing to their orthogonality (Fig. 2e, f). The above methods are well compatible with brain tissue. We immunostained mouse brain slices for presynaptic and postsynaptic markers Bassoon and Homer1 using secondary antibodies conjugated with NHS-MA-biotin and NHS-MA-DIG, and then treated these slices with digestion, post-expansion staining, and expansion (Fig. 2g). Presynaptic and postsynaptic densities were well separated and junctions between them were clearly observable with a confocal microscope (Fig. 2 h-j).

**Fig. 2.**
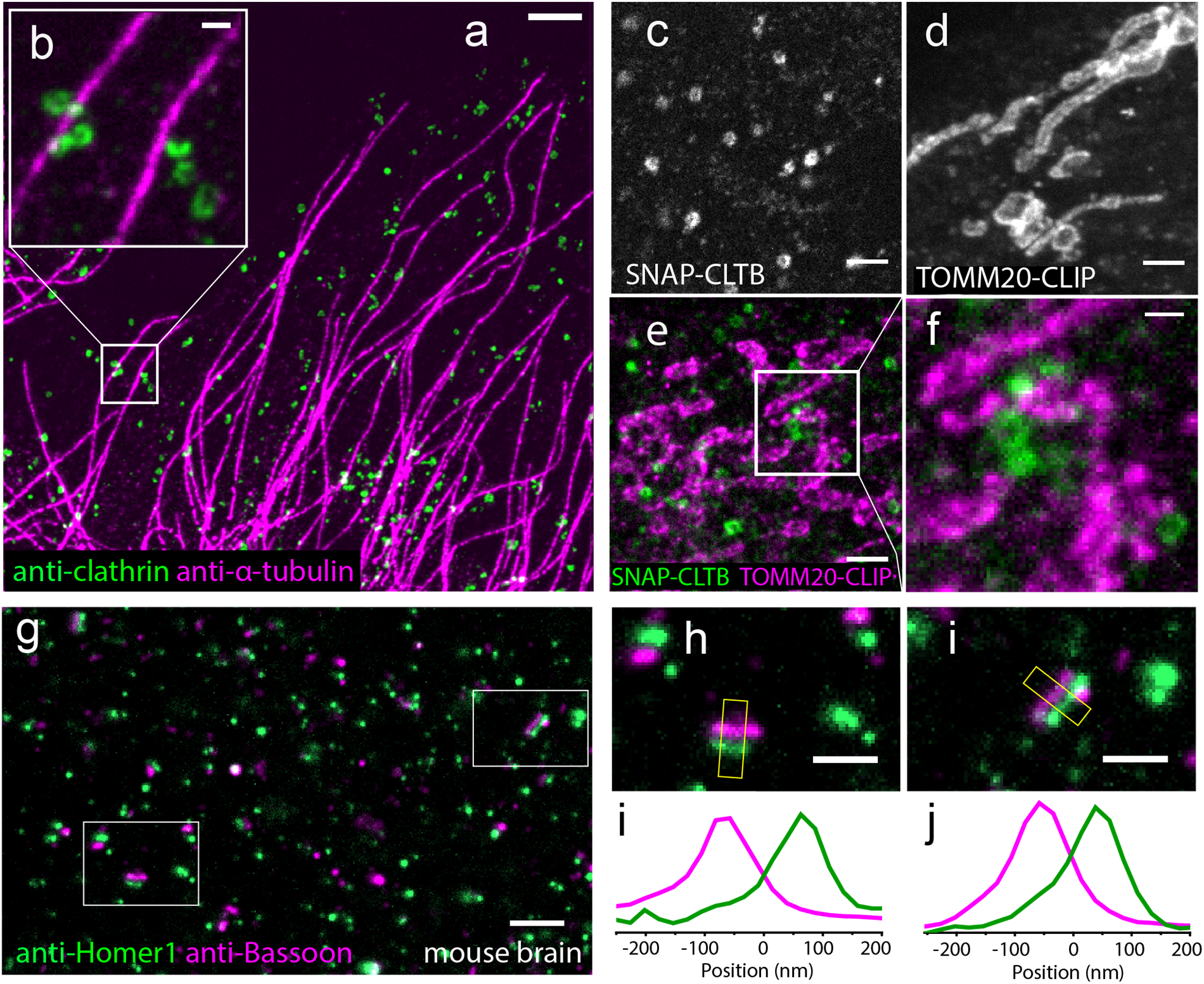
Two color LR-ExM images using immunostaining and protein-tag approaches. (a) Two color LR-ExM confocal image of microtubules labeled with NHS-MA-Biotin conjugated secondary antibodies (magenta) and CCPs labeled with NHS-MA-DIG conjugated secondary antibodies (green) in a U2OS cell, with a magnified view (b). (c-f) LR-ExM confocal images of CCPs and/or mitochondria in HeLa cells labeled using (c) SNAP tag labeled clathrin, (d) CLIP tag labeled TOMM20, and (e) two-color imaging with a magnified view (f). (g) LR-ExM confocal image of mouse brain slice indirectly immunostained for the presynaptic marker Bassoon (magenta) and the postsynaptic marker Homer1 (green), with (h, i) zoomed-in images of synapses and (i, j) transverse intensity profiles along the yellow box long axes. Bassoon is labeled with NHS-MA-DIG conjugated secondary antibodies, and Homer1 is labeled with NHS-MA-Biotin conjugated secondary antibodies. All samples are post-expansion stained with streptavindin-AF488 and or anti-Digoxin-AF594. The length expansion ratios for images (a/ b), (c), (d), (e/f), and (g/h/i) are 4.7, 4.4, 4.4, 4.5 and 4.2, respectively. Scale bars, 1 µm for (a) & (g), 200 nm for (b), (f), (h) & (i), and 500 nm for (c-e). All scale bars in pre-expansion units.

The enzymatic and immunostaining LR-ExM approaches can be combined easily. As a demonstration, we imaged nuclear lamina with SNAP-tag labeled Lamin A/C (LMNA) and antibody-stained nuclear pore complex (NPC) (Fig. 3a-d). Nuclear lamina is a dense fibrillar network bridging the nuclear envelope and chromatin. It positions the nuclear pore complexes (*32*) and participates in chromatin organization (*33, 34*). Two-color LR-ExM cleanly resolved the holes in the nuclear lamina where NPCs are located. The high labeling efficiency of enzymatic protein tags was the key to achieving superior image quality compared to previous super-resolution microscopy results using antibody staining (*35*). The area of the holes in the Lamin A/C network varies from 0.1 to 0.5 µm^2^, with an average of 0.35 µm^2^ (Fig. 3e), which is in agreement with previous SIM studies (*35*) and electron microscopy (EM) (*32*). We observed a strong anti-correlation between Lamin A/C and NPC signal (Fig. 3a, b).

**Fig. 3.**
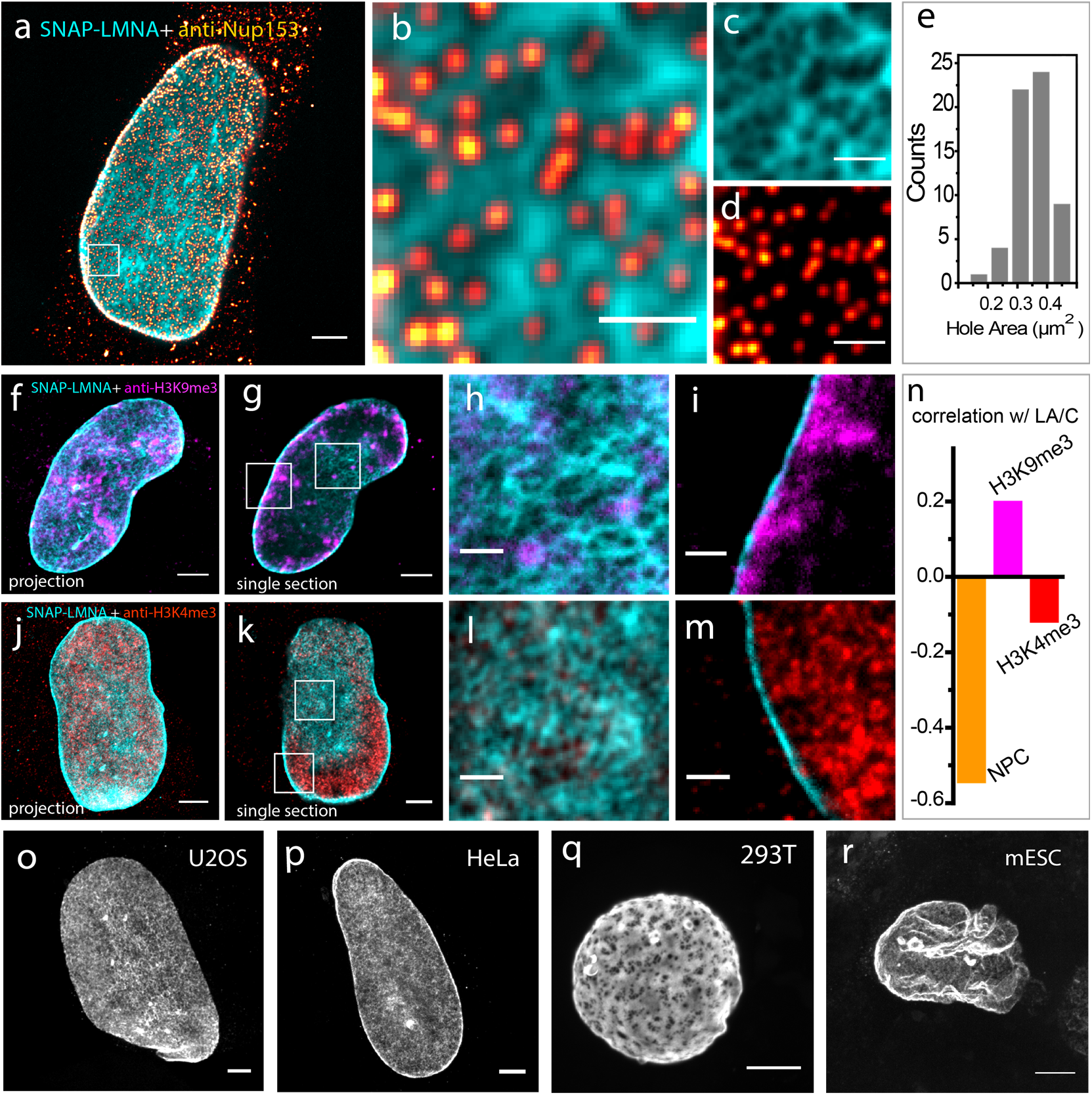
LR-ExM reveals subcellular protein organizations. (a) Two-color confocal LR-ExM of SNAP tagged Lamin A/C (cyan) and immunostained NPC (red hot) of a HeLa cell, with (b) magnified view and (c, d) views of individual channels of (b). Note the cytoplasmic background in (a) is caused by the anti-NUP153 antibody. (e) Histogram of lamin hole area in the boxed region. (f-i) Two color confocal LR-ExM of SNAP tagged Lamin A/C (Cyan) and immunostained H3K9me3 (magenta) of a HeLa cell, with (f) a maximum intensity project of a z stack covering the bottom half of the nucleus, (g) a single section of the nucleus, and (h & i) magnified views of the boxed regions in (g). (j-m) Two-color confocal LR-ExM of SNAP tagged Lamin A/C (Cyan) and immunostained H3K4me3 (red), with (j) a maximum intensity project of a z stack covering the bottom half of the nucleus, (k) a single section of the nucleus, and (l & m) magnified views of the boxed regions (k). (n) The correlation coefficients of NPC with Lamin A/C, H3K9me3 with Lamin A/C, and H3K4me3 with Lamin A/C. (o-r) Confocal LR-ExM of Lamin A/C in (o) U2OS, (p) HeLa, (q) HEK 293T, and (r) mESC cells, showing maximum intensity projections over the bottom half of a nucleus. The length expansion ratios for images (a/b/c/d), (f/g/h/i), (j/k/l/m), (o), (p), (q), and (r) are 4.5, 4.5, 4.3, 4.2, 4.3, 4.6 and 4.4, respectively. Scale bars: 2 µm (a, f, g, j, k, & o-r), and 500 nm (b-d, h, i, l, & m). All scale bars in pre-expansion units.

We further characterized the spatial relationship between Lamin A/C network and two different chromatin markers: H3K9me3 for repressed chromatin (*36*) (Fig. 3c-i) and H3K4me3 for active chromatin (*37*) (Fig. 3d-m). With the enhanced spatial resolution, two-color LR-ExM images clearly showed that, near the nuclear envelope, H3K9me3 was concentrated near Lamin A/C-rich regions, whereas H3K4me3 had an anti-correlation with Lamin A/C signal in a similar manner as NPC (Fig. 3n). This result agrees with the model for the lamin-association of heterochromatin and NPC-association of euchromatin at the nuclear periphery. We were also able to resolve the distinct fine network features of Lamin A/C in a variety of cell lines including mouse embryonic stem cells (Fig. 3o-r). All of these results illustrate the application of LR-ExM (potentially in conjugation with fluorescence *in situ* hybridization to visualize DNA) in studying chromatin organization and sub-diffraction-limit-sized chromatin domains such as lamin-associated domains.

The high label retention of trifunctional anchors and high labeling efficiency of enzymatic tags in LR-ExM enhances its combination with super-resolution microscopy. We first demonstrated this application by imaging antibody-stained distal appendages in primary cilia of Retinal Pigment Epithelium (RPE) cells using LR-ExM combined with SIM (LR-ExSIM) (Fig. 4a). The LR-ExSIM image clearly showed 9 clusters of distal appendage marker CEP164, achieving similar resolution and image quality to the STORM image of the same target protein (Fig. 4b). The symmetry and size of distal appendages measured using LR-ExM agreed with the model based on STED (*38*) and STORM studies (*39*) (Fig. 2c). We calculated the resolution (FWHM) of LR-ExSIM to be 34 nm.

**Fig. 4.**
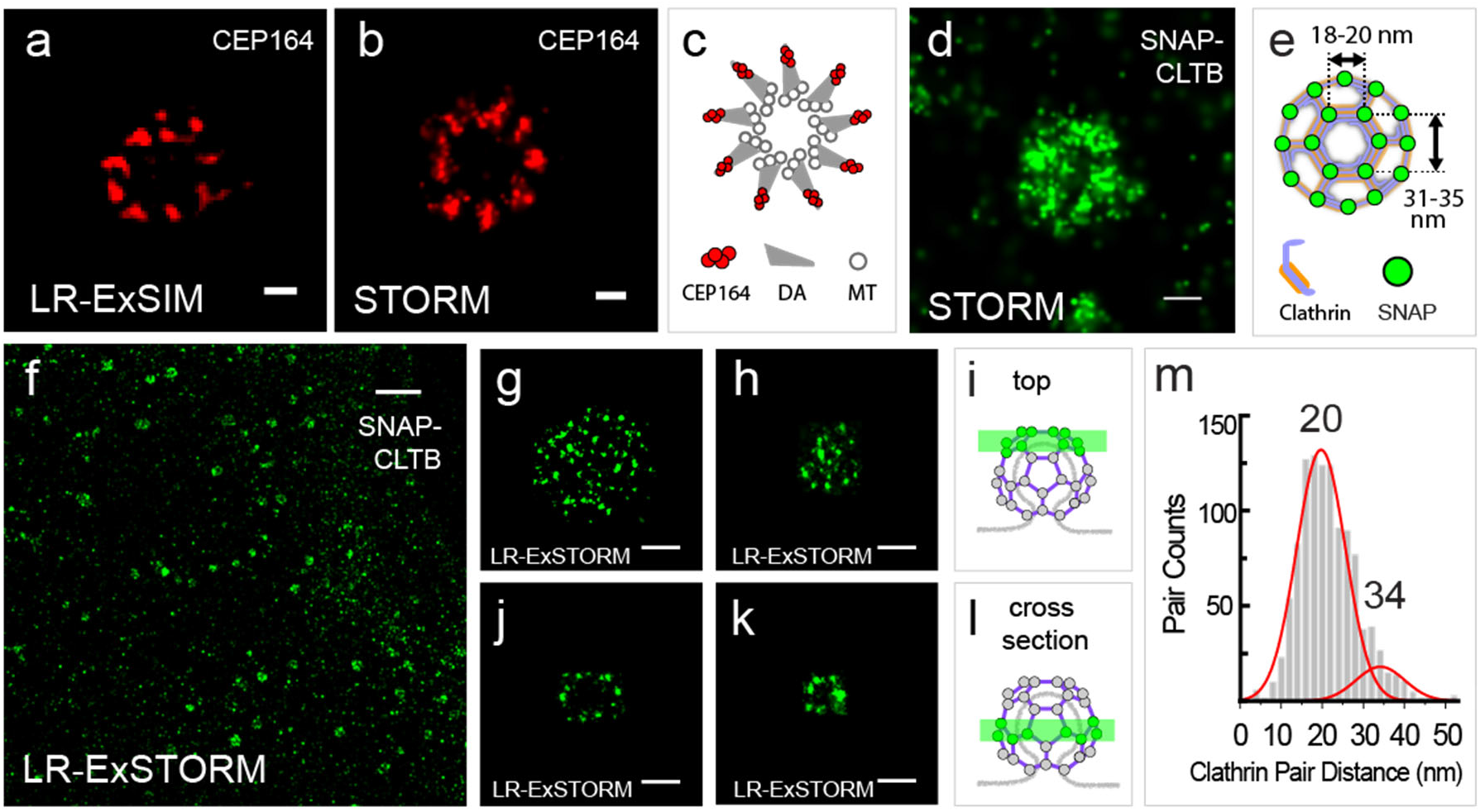
LR-ExSIM and LR-ExSTORM reveals subcellular protein organizations. (a) LR-ExSIM image of Cep164 in distal appendages of a primary cilium indirectly immunostained with NHS-MA-Biotin secondary antibodies, comparing with (b) STORM image of Cep164 distal appendages of un-expanded cilium. (c) Schematic of the structure of distal appendages of the primary cilium. (d) STORM image of an un-expanded HeLa cell over expressing SNAP-CLTB, stained with BG-AF647. (e) Schematic of the structure of a CCP with SNAP-tag-labeled CLTB. (f) LR-ExSTORM image of a HeLa cell over expressing SNAP-CLTB, stained with BG-MA-biotin, and post-expansion labeled with streptavidin-AF647. (g & h) Images of x-y cross sections at the top of single CCPs as illustrated in (i). (j & k) Images of x-y cross sections in the middle of singles CCPs as illustrated in (l). (g-k) are different CCPs. (m) Nearest cluster distance analysis of 134 CCPs imaged with LR-STORM. The length expansion ratios for images (a), (f), (g), (h), (j), and (k) are 4.2, 3.3, 3.3, 3.1, 3.1, and 3.1, respectively. The length expansion ratio for samples used in plot (m) is 3.2 ± 0.2. Scale bars: 100 nm for (a, b, g-k), 200 nm for (d), 2 µm for (f). All scale bars in LR-ExM images are in pre-expansion units.

Finally, we pushed forward the resolution to the molecular level by combining LR-ExM with STORM (LR-ExSTORM). We have examined commonly used photoswitchable dyes including Cy5, Cy5.5, and AF647, all of which show no loss in either molecular brightness or photoswitching kinetics compared to non-expansion STORM conditions (*30*). The length expansion ratio for LR-ExSTORM ranges from 3 to 3.3. It is smaller than 4 because the STORM mounting medium for dye photoswitching has a relatively high ionic strength. In U2OS cells expressing SNAP-tag labeled clathrin light chain B (CLTB), we demonstrated that LR-ExSTORM revealed far more details than STORM alone (Fig. 4d-m). While the STORM showed the hollow structure of the CCP (Fig. 4d), The LR-ExSTORM was able to resolve the lattice vertices of CCPs as clusters of localization points, each cluster resulting from repetitive photoswitching of one AF647 label (Fig. 4g-k). The effective localization precision of LR-ExSTORM was measured to be ∼ 4 nm (FHWM of clusters rescaled by the expansion ratio), which is comparable to the size of a typical protein molecule.

At such a small scale, localization precision is not the only parameter to determine image resolution. The distortion of hydrogel at nanoscale and needs to be considered. In addition, the size and orientation of the label also add uncertainty to measurement. To evaluate the uncertainty, we compared the unit length of clathrin lattice measured by ExSTORM and EM. In ExSTORM images focused at the top of CCPs where the lattice plane is horizontal (e.g. Fig. 4g, h), we measured distances from the centroid of one cluster to the centroid of its nearest neighbor. The histogram of these nearest neighbor distances (1102 pairs from 134 CCPs) showed a main peak at 20 nm and a small shoulder peak at 34 nm (Fig. 4m). The main peak matched the distance between adjacent vertices of clathrin lattice as previously measured by EM (*40, 41*), while the shoulder peak matched the distance between every other vertex (Fig. 4e). This agreement confirmed the ability of LR-ExSTORM to faithfully expand protein complexes at the 10-20 nm scale, possibly attributed to the high degree of isotropic expansion. The 6 nm standard deviation of the histogram is contributed by polyacrylamide gel distortion, the SNAP tag, and the heterogeneity in the clathrin lattice. Moreover, the histogram also indicates that our labeling efficiency has resulted in a majority of vertices containing at least one labeled CLTB, noting that not all clathrin light chains had SNAP-tag in our case because of the presence of endogenous CLTB and the other clathrin light chain isoform, CLTA.

## DISCUSSION

We developed LR-ExM that eliminates the fluorescence loss caused by both polymerization and digestion using trifunctional anchors, and therefore improved the fluorescence intensity by several times. Paring the trifunctional anchors with SNAP and CLIP tags, LR-ExM achieved high labeling efficiency. It paved the way for the combination of ExM and super-resolution microscopy, which elevates the resolution of optical microscopy to the molecular scale. For such high resolution (<10 nm), high labeling efficiency of individual target proteins rather than overall brightness is required to obtain complete molecular architecture of protein complexes. This requirement is much more stringent, compared with diffraction limited imaging. The high labeling efficiency is a major advantage of LR-ExM in combining with super-resolution microscopy, such as STORM, Photoactivated Localization Microscopy (PALM), Stimulated Emission Depletion (STED) microscopy and SIM.

There are other methods that can efficiently recover the fluorescent signal lost in polymerization reactions. For example, Immunosignal hybridization chain reaction (isHCR) *(22)* and immunostaining with signal amplification by exchange reaction using hybridization chain reaction (Immuno-SABER) *(23)* increase fluorescence by two to three orders of magnitudes, which remarkably boosts the detection sensitivity of tissue imaging. However, these methods are unable to rescue the lost information from antibodies that are washed away during the digestion step, resulting in incompletely but still brightly labeled structures. Moreover, the bulky labels will introduce localization displacement that is detectable with the molecular resolution. Compared with these label amplification methods, LR-ExM not only avoids the polymerization-induced fluorophore damage by post-expansion labeling, but eliminates the digestion-induced fluorophore loss by directly crosslinking the reporter to the hydrogel. Wen and coworkers presented a similar chemical linking approach that enables direct grafting of a targeting molecule and fluorophore to the hydrogel, and demonstrated the design principle of trifunctional anchors also work for antibody-free staining of small biomolecules like lipid and actin in ExM (*42*).

To address the limitations of immnunostaining, we developed the enzymatic approach of LR-ExM using SNAP, CLIP tags, and trifunctional anchors bearing BG or BC as the connector group. This approach is ideal for molecular-resolution microscopy with the advantages of the high ligand-binding efficiency of SNAP or CLIP tag, and their much smaller size than antibodies (19 kDa SNAP vs. 150 kDa IgG). For the same reasons, cell lines with SNAP tagged nuclear pore protein are used to benchmark the quality of super-resolution microscopes (*19*). Compared with diffraction limited imaging, molecular-resolution imaging has more stringent requirement for small labels. LR-ExM minimizes the size of the label by using not only small enzymatic tags, but also short trifunctional anchors only about 1 nm long. We demonstrated the resolving power of LR-ExSTORM by visualizing the polyhedral lattice of CCPs *in situ* (Fig. 4 f-m), which is not resolvable by using STORM alone (Fig. 4d) (*43, 44*). While using SNAP and CLIP tags can improve the resolution by reducing the tag size and can provide high labeling efficiency, it does require genetic approaches to fuse the tags to the targeted proteins. Extra attention needs to be paid to the expression level of enzymatic tags, as over expression may cause structural or functional artifact. We recommend to keep the expression level similar to the endogenous level of the target protein.

Another advantage of the LR-ExM is the robustness. Most ExM protocols consist of nonspecific protein-hydrogel anchoring with chemical crosslinkers (e.g. MA-NHS, and glutaraldehyde) and proteinase digestion (*2, 3, 5, 6, 8-14, 22, 23*). How much fluorescence is retained depends on the balance of anchoring density and digestion efficiency, resulting in largely variable fluorescence intensity across experiments (*2, 4, 18*). Higher anchoring density increases fluorescence retention, while stronger digestion duration decreases fluorescence retention. However, the fluorescence retention rate of LR-ExM is independent from digestion conditions, since the reporter is directly anchored to the hydrogel, not through antibodies. Consequently, LR-ExM protocol is robust and reproducible across experiments.

LR-ExM is also a versatile method that can be integrated with ExM protocols targeting nucleic acids (e.g. ExFISH (*15*)), lipid (e.g. mExM(*45*), TRITON(*42*)), context protein (e.g. FLARE(*46*), pan-ExM(*7*)), and so on. Although only two-color LR-ExM using biotin-and digoxingenin-bearing trifunctional anchors was demonstrated, additional color channels were imaged by post-expansion labeling DNA with DAPI, and post-expansion labeling telomere with LNA oligonucleotides (data not shown). The multiplexity can be further extended by developing new trifunctional anchors with more small chemical reporters, such as alkyne (pairing with azide) and chloroalkane (pairing with HaloTag).

In summary, LR-ExM is an effective, robust, and versatile method to enhance the signal and labeling efficiency of expansion microscopy. Our trifunctional anchors can be applied to both antibody and enzymatic labeling. They can also be integrated into most existing ExM protocols, greatly increasing their signals and multiplexity. Overcoming the bottleneck of label loss, the currently achieved post-expansion resolutions of ∼70 nm with confocal microscopy, ∼30 nm with SIM, and localization precision of ∼4 nm with STORM are suitable to cover a wide range of biological questions at various scales.

## MATERIALS AND METHODS

### Trifunctional anchor synthesis

We synthesized five trifunctional anchors, including HOOC-MA-Biotin, HOOC-MA-DIG, BG-MA-Biotin, BG-MA-DIG, and BC-MA-DIG (Fig. S1). HOOC-MA-Biotin and HOOC-MA-DIG anchors were converted to NHS-MA-Biotin and NHS-MA-DIG respectively to conjugate antibodies for the immunostaining approach of LR-ExM. The BG-MA-Biotin, BG-MA-DIG, and BC-MA-DIG anchors were directly used for the protein-tag approach of LR-ExM. The synthetic schemes are shown in Fig. S2.

All reactions were performed in flame- or oven-dried glassware fitted with rubber septa under a positive pressure of nitrogen, unless otherwise noted. All reaction mixtures were stirred throughout the course of each procedure using Teflon-coated magnetic stir bars. Air- and moisture-sensitive liquids were transferred via syringe. Solutions were concentrated by rotary evaporation below 30 °C. Analytical thin-layer chromatography (TLC) was performed using glass plates pre-coated with silica gel (0.25-mm, 60-Å pore size, 230−400 mesh, SILICYCLE INC) impregnated with a fluorescent indicator (254 nm). TLC plates were visualized by exposure to ultraviolet light (UV) and then were stained by submersion in a basic aqueous solution of potassium permanganate or with an acidic ethanolic solution of anisaldehyde, followed by brief heating. For synthetic procedures and characterization data (thin-layer chromatography (TLC), NMR and Mass Spectroscopy), see Supplementary Methods.

### Antibodies

The following primary antibodies were used for the immunostaining approach of LR-ExM: rabbit anti-Clathrin heavy chain antibody (Abcam, ab21679), rat anti-alpha-Tubulin Antibody, tyrosinated, clone YL1/2 (Millipore Sigma, MAB1864-I), and monoclonal mouse anti-Nup153 antibody (Abcam, ab24700), anti-H3K9me3 (Abcam, ab176916), anti-H3K4me3 (Abcam, ab8580). The secondary antibodies used for trifunctional anchor conjugation are: Unconjugated AffiniPure Donkey Anti-Rabbit IgG (H+L) (Jackson ImmunoResearch 711-005-152), and Unconjugated AffiniPure Donkey Anti-Rat IgG (H+L) (Jackson ImmunoResearch 712-005-153).

### Antibody conjugation

Secondary antibodies were labeled with the amine-reactive trifunctional anchor: NHS-MA-Biotin or NHS-MA-DIG. Amine-reactive trifunctional anchors were freshly made from stock solutions of synthesized trifunctional anchors HOOC-MA-Biotin or HOOC-MA-DIG (26 mM, in DMSO, stored at -20 °C). EDC solution (20 µL, 100 mg mL^-1^), NHS solution (20 µL, 65 mg mL^-1^) and DMAP solution (10 µL, 30 mg mL^-1^) were sequentially added into the solution of HOOC-MA-Biotin or HOOC-MA-DIG (50 μL, 26 mM). The mixture was gently shaken at room temperature for 16 hours while shielding from light with aluminum foil. Using the aforementioned volumes, the final concentration of in situ prepared NHS-MA-Biotin or NHS-MA-DIG is 13 mM.

To conjugate the secondary antibodies with the amine-reactive trifunctional anchor, the following procedure was performed: 10 µL aqueous NaHCO_3_ solution (1 M) was added to an Eppendorf tube containing 80 µL of unconjugated antibody solution (1mg mL^-1^). A solution of the amine-reactive trifunctional anchor (NHS-MA-Biotin or NHS-MA-DIG, 13 mM, 24 µL) was then added to the NaHCO_3_-buffered antibody solution. The labeling reaction mixture was gently rocked for 20 min at room temperature. During the reaction, a Sephadex G-25 column (GE Healthcare, NAP-5) was equilibrated with PBS (PH 7.4). The labeling reaction mixture was loaded onto the column, followed by flowing with 650 μL of PBS. The antibodies conjugated with trifunctional anchors were collected by eluting the column with another 250 μL of PBS, and stored at 4 °C.

The procedure of antibody conjugation with commercially available bifunctional linker NHS-Biotin was the same to the conjugation with trifunctional anchors, except that a solution of NHS-Biotin (26 mM, 4 µL) instead of the trifunctional anchor was added to the NaHCO_3_-buffered antibody solution.

### Quantification of biotin-to-antibody ratio

Antibody concentration was characterized by measuring the absorption at 280 nm with a UV-Vis spectrophotometer. The concentration of biotin was measured using HABA/Avidin reagent kit, following the protocol provided by the supplier (Thermo Scientific™ Pierce™ Biotin Quantitation Kit, # 28005). The biotin-to-antibody ratios of the antibody conjugated with NHS-Biotin in Fig. 1d (Fig. S3b), the antibody conjugated with NHS-MA-Biotin in Fig. 1e (Fig. S3d), and the antibody conjugated with NHS-Biotin and NHS-MA in Fig. S3c are 12.1, 8.6, and 9.9, respectively. The dye-to-antibody ratio of the antibody conjugated with Alexa Fluor 488 in Fig. 1c, (Fig. S3a) is 8.9. These biotin-to-antibody ratios and the dye-to-antibody ratio are used to normalize the label retention efficiency of proExM, biotin-ExM, and LR-ExM (Fig. 1f, Fig. S3i).

### Cell culture

U2OS cells were cultured in McCoy’s 5a (ATCC, 30-2007) supplemented with 10% FBS at 37 °C in 5% CO_2_. HeLa, and HEK 293T cells were cultured in DMEM-Glutamax (Thermo Fisher) supplemented with 10% FBS at 37 °C in 5% CO_2_. U2OS, HeLa, and HEK 293T cells were seeded at 1 × 10^4^ cells/cm^2^ in 16-well chambers (Grace Bio-Labs, 112358) and grown to 75% confluency.

Human retinal epithelial (RPE1-hTERT) cells were cultured in DMEM/F12 supplemented with 10 % FBS at 37 °C in 5% CO_2_. Cells were plated on 16-well chambered slides coated with 0.1% gelatin (G1393 from Sigma) at 1 × 10^4^ cells/cm^2^ per well, and serum starved in OptiMEM reduced serum media for 24 h to induce ciliation. Cell lines were not authenticated. No commonly misidentified cell lines were used. All growing cell lines were routinely tested for mycoplasma.

### Molecular cloning

To generate the pTOMM20-N-10-CLIPf mammalian expression plasmid, the DNA of CLIPtag was PCR amplified from pCLIPf vector (plasmid source: the Michael Davidson Fluorescent Protein Collection at the UCSF Nikon Imaging Center) using primers (Forward: GCGGGGATCCACCGGTCGCCACCATGGACAAAGACTGCGAAATGAAGC. Reverse: TCTAGAGTCGCGGCCGCTTAACCCAGCCCAGGCTTGCCC). We then performed restriction enzyme digestion on vector pmEmerald-TOMM20-N-10 (plasmid source: the Michael Davidson Fluorescent Protein Collection at the UCSF Nikon Imaging Center): cutting out the mEmerald sequence between BamHI and NotI. The PCR amplified CLIPtag were then ligated with the digested vectors using In-Fusion HD Cloning kit (Clontech). The plasmids pSNAPf-Clathrin-15 and pSNAPf-LMNA were directly purchased from UCSF Nikon Imaging Center. For constructing the lentiviral production vectors, DNAs of TOMM20-N-10-CLIPf and SNAPf-Clathrin-15 were directly PCR amplified from mammalian expression constructs and subcloned into lentiviral pHR-SFFV vector (BamHI/NotI) using In-Fusion HD Cloning kit (Clontech).

### Immunostaining

For microtubule immunostaining and microtubule-clathrin co-immunostaining, the cells were fixed with 3.2% PFA in PEM buffer (100 mM PIPES, 1 mM EGTA, 1 mM MgCl2, pH 6.9) at room temperature for 10 min. The fixation was reduced with 0.1% sodium borohydride in PBS for 7 min. The sodium borohydride was removed by washing with PBS three times with 5 minutes of incubation between washes. The fixed cells were incubated in blocking buffer (PBS, 3% BSA, 0.1% Triton X-100) for 30 minutes at room temperature. Primary antibodies at a concentration of 2 μg ml^-1^ were added to the fixed cells in blocking buffer for 16 h at 4 °C. The primary antibodies used or this paper are listed in the Supplementary Information. After, primary antibody incubation, the cells were washed with PBS three times with 5 minutes of incubation between washes. Secondary antibodies conjugated with trifunctional anchors were added at a concentration of 3 μg ml^-1^ and incubated for 1 hour in blocking buffer on an orbital shaker. The secondary antibodies were removed by three washes with PBS buffer.

For CEP164 immunostaining, cells were fixed with 100% cold methanol for 3 minutes, and then incubated in blocking buffer (2.5% BSA, 0.1% Triton-X 100 in PBS) for 1 hour at room temperature and incubated with primary antibody in blocking buffer (1:100 dilution of goat anti-CEP164, sc-240226 from Santa Cruz biotechnology) overnight at 4 °C. Cells were washed with PBS four times with 5 minutes of incubation between washes, and incubated with donkey anti-goat secondary antibody conjugated with NHS-MA-Biotin anchors for 1 h at room temperature. The secondary antibodies were removed by three washes with PBS buffer.

For mouse brain tissue immunostaining, wild-type adult mouse brain was fixed in 4% paraformaldehyde overnight, before transferring to 30% sucrose and then OCT for cryoprotection. Tissue block was sectioned coronally at 20 μm thick and incubated in blocking buffer (10% goat serum, 3% BSA, 1% glycine, 0.4% Triton-X 100 in TBS) for 1 hour at room temperature. Tissue slices were then stained with primary antibodies (1:500 dilution of mouse anti-Bassoon, VAM-PS003 from StressGene; 1:200 dilution of rabbit anti-Homer1, 160003 from Synaptic Systems) in blocking buffer overnight at room temperature. Slices were washed with TBS three times for 5 minutes each time and then incubated with donkey anti-mouse (conjugated with NHS-MA-DIG) and donkey anti-rabbit (conjugated with NHS-MA-Biotin) secondary antibodies (both at 1:100 dilution in blocking buffer) for 2 hours at room temperature.

### SNAP- and CLIP-tag labeling

The cells expressing SNAP-tag and or CLIP-tag were fixed for 10 min with 4% PFA in PBS buffer. The PFA was removed by PBS washing. The fixed cells were incubated in blocking buffer (PBS, 3% BSA, 0.1% Triton X-100) for 30 minutes at room temperature. The cells were then incubated in 3 μM trifunctional anchor SNAP-MA-Biotin and or 5 μM CLIP-MA-DIG for 1 h.

### Polymerization, proteinase digestion, post-expansion labeling, and expansion

The polymerization and proteinase digestion steps are similar to the proExM protocol (*2, 3*) with two exceptions. One is that we also treated the sample with DNAse I prior to plolemerization to fragment the genomic DNA, with the intention to reduce potential distortions inside and around the nucleus. The other exception is that protein anchoring with methacrylamide NHS ester or glutaraldehyde is not necessary but optional.

Specifically, fixed cells were incubated in DNAse I buffer (New Englab BioLabs, M0303S, 1:100 dilution in PBS buffer) for 30 min at 37 °C, and then were polymerized in a mixture of monomer solution (8.6 g Sodium acrylate, 2.5 g Acrylamide, 0.15 g N,N′-Methylenebisacrylamide, 11.7 g Sodium chloride per 100 mL PBS buffer), TEMED (final concentration 0.2% (w/w)) and ammonium persulfate (final concentration 0.2% (w/w)) for 1 h at 37 °C. The gel was then digested with proteinase K (Sigma-Aldrich, P4850-5ML) with the final concentration of 8 units mL^-1^ in digestion buffer (50 mM Tris PH 8.0, 1 mM EDTA, 0.5% Triton X-100, 0.8 M guanidine HCl) for 18 h at room temperature or 4 h at 37 °C. After digestion, the proteinase K was removed by four washes with excessive water, for 30 min each time. To introduce fluorophores to the trifunctional anchors on the target cellular structures, we incubated the hydrogel in 2 μg mL^-1^ Alexa Fluor 488 labeled Streptavidin (Jackson ImmunoResearch Laboratories, 0165400084) and or DyLight 594 Labeled Anti-Digoxigenin/Digoxin (DIG) (Vector Laboratories, DI-7594) in HEPES buffer (10 mM HEPES, 15m mM NaCl, pH 7.5) for 24 h. For LR-ExSTORM and LR-SIM, Alexa Fluor 647 Streptavidin (Jackson ImmunoResearch Laboratories, 0160600084) was used for post-expansion staining. The post-expansion labeled hydrogel was then washed and expanded by four washes with excessive water, at least 30 min each time. It is optional to treat the cells with 25 mM methacrylic acid N-hydroxysuccinimide ester for 1 h before polymeration.

The length expansion ratio of the LR-ExM protocol is determined by measuring the diameters of the gel before and after expansion with calipers.

### STORM image acquisition and analysis

STORM was performed on a custom-built microscope based on a Nikon Ti-U inverted microscope. Two lasers (Coherent CUBE 405 and CUBE 642) were combined using dichroic mirrors, aligned, expanded and focused to the back focal plane of the objective (Nikon Plan Apo 100x oil NA 1.45). The lasers were controlled directly by the computer. A quad band dichroic mirror (zt405/488/561/640rpc, Chroma) and a band-pass filter (ET705/70m, Chroma) separated the fluorescence emission from the excitation light. During image acquisition, the focusing of the sample was stabilized by a closed-loop system that monitored the back reflection from the sample coverglass via an infrared laser beam sent through the edge of the microscope objective. A low-end piezoelectric deformable mirror (DM) (DMP40-P01, Thorlabs) was added in the emission path at the conjugate plane of the objective pupil plane^28^. By first flattening the mirror and then manually adjusting key Zernike polynomials, this DM corrected aberrations induced by both the optical system and the glass-water refractive index mismatch when the sample was several micrometers away from the coverglass. The fluorescence was recorded at a frame rate of 57 Hz on an electron multiplying CCD camera (iXon+ DU897E-CS0-BV, Andor).

The mounting medium used for STORM imaging was water with the addition of 10mM mercaptoethylamine at pH 8.5, 5% glucose (w/v) and oxygen scavenging enzymes 0.5 mg/ml glucose oxidase (Sigma-Aldrich), and 40 mg/ml catalase (Roche Applied Science). The pH of the final solution is adjusted to 8.4. The mounting medium remained suitable for imaging for 1**–**2 h. Photoswitchable dye Alexa Fluor 647 was conjugated on streptavidin and was used for imaging with a ratio of 0.8 to 1 dye per streptavidin. The stained hydrogel was incubated in the mounting medium for 15 min before mounting. The hydrogel was them transferred to laser-cut sample chamber with a polylysine-coated coverglass that we devised to mechanically stabilize the expanded gel during image acquisition (Fig. S4).

Alexa Fluor 647 was excited with a 642 nm imaging laser, with typically 1 kW cm^-2^ laser intensity at the focal plane. Analysis of STORM raw data was performed in the Insight3 software^28^, which identified and fitted single molecule spots in each camera frame to determine their **x** and **y** coordinates as well as photon numbers.

### Drift reduction and correction

We minimized the sample drift during data acquisition by mounting the hydrogel in a 3D-printed chamber (Fig. S4). The bottom of the chamber is a coverglass modified with poly-L-lysine, which creates a strong adhesion to the negatively charged hydrogel. The drift during data acquisition was further corrected using imaging correlation analysis. The drift-corrected coordinates, photon number, and the frame of appearance of each identified molecule were saved in a molecule list for further analysis.

### Storage and reimaging

The imaged hydrogel can be stored in PBS buffer. The fluorescence will retain for at least a month for multiple rounds of imaging. Note that in water, fluorescence signal will completely fade away in a week.

### Quantification and comparison of fluorescence retention efficiencies

To compare the fluorescence retention efficiencies of different ExM methods, we took the images of immunostained microtubules in U2OS cells prepared with different ExM methods, with the same imaging condition. We calculated the retained fluorescence by dividing the total fluorescence intensity of the all microtubules by their total length. The total length of all the microtubules in each image was quantified using a Fiji plugin JFilament. The quantification process and results are shown in Fig. S3.

### Quantification of LR-ExSTORM images of CCPs

We LR-ExSTORM imaged CCPs in U2OS cells expressing SNAP-tag labeled clathrin light chain B (CLTB) stained with BG-MA-biotin anchor before expansion and biotin-Alexa Fluor 488 after expansion. CCPs focused at the top were selected for the quantification. We measured the distances from the centroid of one cluster to the centroid of its nearest neighbor in the central area of each CCP, and excluded the clusters at the CCP edge to avoid off-focus localizations. We plotted the histogram of these nearest neighbor distances (1102 pairs from 134 CCPs), and fitted the distance distribution with Gaussian functions. The position of the center and the standard deviation of the gaussian peak are respectively used as the distance between neighboring vertices of the polyhedral CCPs and the standard deviation of the distance (Fig. 2). The effective localization precision of LR-ExSTORM was calculated by dividing the mean of measured FHWM of clusters in the STORM images by the length expansion ratio. Each cluster is the superimposition of fitted Gaussian peaks of repetitive photoswitching of one AF647 label.

### Trifunctional anchor availability

Samples of trifunctional anchors described in this manuscript are available on request.

## SUPPLEMENTARY MATERIALS

### Materials and Methods

Fig. S1. Structures of trifunctional anchors.

Fig. S2. Synthetic scheme of trifunctional anchors.

Fig. S3. Comparison of fluorescence intensities resulting from different ExM methods.

Fig. S4. 3D-printed chamber for drift reduction of hydrogel.

Fig. S5. LR-ExSIM of microtubules.

Fig. S6. Resolution measurement for LR-ExM confocal images.

Fig. S7. LR-ExSIM and STORM reveal structure of distal appendages with similar super resolution.

## ACKNOWLEDGEMENTS

We thank Drs. Ed Boyden and Fei Chen for their help with the proExM protocol, and Drs. Joshua C Vaughan and Aaron Halpern for the help with their ExM protocol. We are grateful to Dr. Keith DeLuca from Jenifer DeLuca Lab for validating the expansion factor of compact structures in the nucleus, Stefan Niekamp from Ron Val Lab for measuring the labeling efficiency, and Dr. Dan Xie for optimizing the deformable mirror, and 3D printing slide adapters for the STORM microscope. We appreciate Dr. Luke Lavis for his consultation on protein tags, Dr. Xiangpeng Li from Adam Abate Lab, and Dr. Xiao Huang from Tejal Desai Lab for their consultation on biochemistry and microfluidics. We thank Eric Simental for transfecting mESC cells. This work was also highly inspired by personal conversations with Drs. David Brown and Juan Guan, and discussions with all the other Huang lab members. This project is supported by the National Institutes of Health (NIH) Director’s New Innovator Award DP2OD008479 and R01GM124334 to B.H., by the NIH Pathway to Independence Award K99GM126136/R00GM126136 and the UCSF Mary Anne Koda-Kimble Seed Award for Innovation to X.S., by the AHA Predoctoral Fellowship 19PRE3480616 to J.C., and by National Science Foundation for a Graduate Research Fellowship 1650113 to A.A.T.. B. H. and J.F.R. are Chan Zuckerberg Biohub Investigators.

## AUTHOR CONTRIBUTIONS

X.S., Q.L., Z.D., I.B.S., and B.H. designed the experiments and interpreted the results. X.S. designed LR-ExM protocols. X.S., Z.L., X.W. and T.C. imaged samples. Q.L. and A.A.T. synthesized the trifunctional anchors, Z.D. did image quantification and antibodies conjugation. S.F. and A.D.R. designed and made plasmids. J.C. prepared and stained the mouse brain. D.K., X.S., J.F.R. and B.H. designed the experiments on imaging primary cilia with ExSIM. D.K. and X.S. took the ExSIM images. X.S. drafted the manuscript. B.H., I.B.S., A.A.T., Z.D., S.F., J.C., E.J.H, J.F.R, and X.S. edited the manuscript.

## DATA AVAILABILITY

All data needed to evaluate the conclusions in the paper are present in the paper and/or the Supplementary Materials.

## REFERENCES

1. F. Chen, P. W. Tillberg, E. S. Boyden, Optical imaging. Expansion microscopy. Science 347, 543–548 (2015).

2. T. J. Chozinski, A. R. Halpern, H. Okawa, H. J. Kim, G. J. Tremel, R. O. Wong, J. C. Vaughan, Expansion microscopy with conventional antibodies and fluorescent proteins. Nat Methods 13, 485–488 (2016).

3. P. W. Tillberg, F. Chen, K. D. Piatkevich, Y. Zhao, C. C. Yu, B. P. English, L. Gao, A. Martorell, H. J. Suk, F. Yoshida, E. M. DeGennaro, D. H. Roossien, G. Gong, U. Seneviratne, S. R. Tannenbaum, R. Desimone, D. Cai, E. S. Boyden, Protein-retention expansion microscopy of cells and tissues labeled using standard fluorescent proteins and antibodies. Nat Biotechnol 34, 987–992 (2016).

4. T. Ku, J. Swaney, J. Y. Park, A. Albanese, E. Murray, J. H. Cho, Y. G. Park, V. Mangena, J. Chen, K. Chung, MAP: Multiplexed and scalable super-resolution imaging of three-dimensional protein localization in size-adjustable tissues. Nat Biotechnol 34, 973–981 (2016).

5. J. B. Chang, F. Chen, Y. G. Yoon, E. E. Jung, H. Babcock, J. S. Kang, S. Asano, H. J. Suk, N. Pak, P. W. Tillberg, A. T. Wassie, D. Cai, E. S. Boyden, Iterative expansion microscopy. Nat Methods 14, 593–599 (2017).

6. S. Truckenbrodt, M. Maidorn, D. Crzan, H. Wildhagen, S. Kabatas, S. O. Rizzoli, X10 expansion microscopy enables 25-nm resolution on conventional microscopes. EMBO Rep 19, e45836 (2018).

7. O. M’Saad, J. Bewersdorf, Light microscopy of proteins in their ultrastructural context. Nat Commun 11, 3850 (2020).

8. C. K. Cahoon, Z. Yu, Y. Wang, F. Guo, J. R. Unruh, B. D. Slaughter, R. S. Hawley, Superresolution expansion microscopy reveals the three-dimensional organization of the Drosophila synaptonemal complex. Proc Natl Acad Sci U S A 114, E6857–E6866 (2017).

9. A. R. Halpern, G. C. M. Alas, T. J. Chozinski, A. R. Paredez, J. C. Vaughan, Hybrid Structured Illumination Expansion Microscopy Reveals Microbial Cytoskeleton Organization. ACS Nano 11, 12677–12686 (2017).

10. Y. Wang, Z. Yu, C. K. Cahoon, T. Parmely, N. Thomas, J. R. Unruh, B. D. Slaughter, R. S. Hawley, Combined expansion microscopy with structured illumination microscopy for analyzing protein complexes. Nat Protoc 13, 1869–1895 (2018).

11. D. Gambarotto, F. U. Zwettler, M. Le Guennec, M. Schmidt-Cernohorska, D. Fortun, S. Borgers, J. Heine, J. G. Schloetel, M. Reuss, M. Unser, E. S. Boyden, M. Sauer, V. Hamel, P. Guichard, Imaging cellular ultrastructures using expansion microscopy (U-ExM). Nat Methods 16, 71–74 (2019).

12. M. Gao, R. Maraspini, O. Beutel, A. Zehtabian, B. Eickholt, A. Honigmann, H. Ewers, Expansion Stimulated Emission Depletion Microscopy (ExSTED). ACS Nano 12, 4178–4185 (2018).

13. R. Li, X. Chen, Z. Lin, Y. Wang, Y. Sun, Expansion enhanced nanoscopy. Nanoscale 10, 17552–17556 (2018).

14. H. Z. Xu, Z. S. Tong, Q. Ye, T. Q. Sun, Z. M. Hong, L. F. Zhang, A. Bortnick, S. Cho, P. Beuzer, J. Axelrod, Q. Z. Hu, M. Wang, S. M. Evans, C. Murre, L. F. Lu, S. Sun, K. D. Corbett, H. Cang, Molecular organization of mammalian meiotic chromosome axis revealed by expansion STORM microscopy. Proceedings of the National Academy of Sciences of the United States of America 116, 18423–18428 (2019).

15. F. Chen, A. T. Wassie, A. J. Cote, A. Sinha, S. Alon, S. Asano, E. R. Daugharthy, J. B. Chang, A. Marblestone, G. M. Church, A. Raj, E. S. Boyden, Nanoscale imaging of RNA with expansion microscopy. Nat Methods 13, 679–684 (2016).

16. K. Chung, J. Wallace, S. Y. Kim, S. Kalyanasundaram, A. S. Andalman, T. J. Davidson, J. J. Mirzabekov, K. A. Zalocusky, J. Mattis, A. K. Denisin, S. Pak, H. Bernstein, C. Ramakrishnan, L. Grosenick, V. Gradinaru, K. Deisseroth, Structural and molecular interrogation of intact biological systems. Nature 497, 332–337 (2013).

17. B. Yang, J. B. Treweek, R. P. Kulkarni, B. E. Deverman, C. K. Chen, E. Lubeck, S. Shah, L. Cai, V. Gradinaru, Single-cell phenotyping within transparent intact tissue through whole-body clearing. Cell 158, 945–958 (2014).

18. S. Truckenbrodt, C. Sommer, S. O. Rizzoli, J. G. Danzl, A practical guide to optimization in X10 expansion microscopy. Nat Protoc 14, 832–863 (2019).

19. J. V. Thevathasan, M. Kahnwald, K. Cieslinski, P. Hoess, S. K. Peneti, M. Reitberger, D. Heid, K. C. Kasuba, S. J. Hoerner, Y. M. Li, Y. L. Wu, M. Mund, U. Matti, P. M. Pereira, R. Henriques, B. Nijmeijer, M. Kueblbeck, V. J. Sabinina, J. Ellenberg, J. Ries, Nuclear pores as versatile reference standards for quantitative superresolution microscopy. Nature Methods 16, 1045–1053 (2019).

20. S. W. Hell, S. J. Sahl, M. Bates, X. Zhuang, R. Heintzmann, M. J. Booth, J. Bewersdorf, G. Shtengel, H. Hess, P. Tinnefeld, A. Honigmann, S. Jakobs, I. Testa, L. Cognet, B. Lounis, H. Ewers, S. J. Davis, C. Eggeling, D. Klenerman, K. I. Willig, G. Vicidomini, M. Castello, A. Diaspro, T. Cordes, The 2015 super-resolution microscopy roadmap. Journal of Physics D: Applied Physics 48, 443001 (2015).

21. D. Kamiyama, B. Huang, Development in the STORM. Developmental Cell 23, 1103–1110 (2012).

22. R. Lin, Q. Feng, P. Li, P. Zhou, R. Wang, Z. Liu, Z. Wang, X. Qi, N. Tang, F. Shao, M. Luo, A hybridization-chain-reaction-based method for amplifying immunosignals. Nat Methods 15, 275–278 (2018).

23. S. K. Saka, Y. Wang, J. Y. Kishi, A. Zhu, Y. Zeng, W. Xie, K. Kirli, C. Yapp, M. Cicconet, B. J. Beliveau, S. W. Lapan, S. Yin, M. Lin, E. S. Boyden, P. S. Kaeser, G. Pihan, G. M. Church, P. Yin, Immuno-SABER enables highly multiplexed and amplified protein imaging in tissues. Nat Biotechnol 37, 1080–1090 (2019).

24. T. C. Murakami, T. Mano, S. Saikawa, S. A. Horiguchi, D. Shigeta, K. Baba, H. Sekiya, Y. Shimizu, K. F. Tanaka, H. Kiyonari, M. Iino, H. Mochizuki, K. Tainaka, H. R. Ueda, A three-dimensional single-cell-resolution whole-brain atlas using CUBIC-X expansion microscopy and tissue clearing. Nat Neurosci 21, 625–637 (2018).

25. Y.-G. Park, C. H. Sohn, R. Chen, M. McCue, D. H. Yun, G. T. Drummond, T. Ku, N. B. Evans, H. C. Oak, W. Trieu, H. Choi, X. Jin, V. Lilascharoen, J. Wang, M. C. Truttmann, H. W. Qi, H. L. Ploegh, T. R. Golub, S.-C. Chen, M. P. Frosch, H. J. Kulik, B. K. Lim, K. Chung, Protection of tissue physicochemical properties using polyfunctional crosslinkers. Nature Biotechnology 37, 73–83 (2018).

26. R. P. J. Nieuwenhuizen, K. A. Lidke, M. Bates, D. L. Puig, D. Grunwald, S. Stallinga, B. Rieger, Measuring image resolution in optical nanoscopy. Nature Methods 10, 557–562 (2013).

27. H. Shroff, C. G. Galbraith, J. A. Galbraith, E. Betzig, Live-cell photoactivated localization microscopy of nanoscale adhesion dynamics. Nature Methods 5, 417–423 (2008).

28. A. Keppler, H. Pick, C. Arrivoli, H. Vogel, K. Johnsson, Labeling of fusion proteins with synthetic fluorophores in live cells. Proceedings of the National Academy of Sciences of the United States of America 101, 9955–9959 (2004).

29. A. Gautier, A. Juillerat, C. Heinis, I. R. Correa, Jr., M. Kindermann, F. Beaufils, K. Johnsson, SNAP-tag: An engineered protein tag for multiprotein labeling in living cells. Chem Biol 15, 128–136 (2008).

30. G. T. Dempsey, J. C. Vaughan, K. H. Chen, M. Bates, X. Zhuang, Evaluation of fluorophores for optimal performance in localization-based super-resolution imaging. Nat Methods 8, 1027–1036 (2011).

31. N. Olivier, D. Keller, P. Gonczy, S. Manley, Resolution doubling in 3D-STORM imaging through improved buffers. PLoS One 8, e69004 (2013).

32. Y. Turgay, M. Eibauer, A. E. Goldman, T. Shimi, M. Khayat, K. Ben-Harush, A. Dubrovsky-Gaupp, K. T. Sapra, R. D. Goldman, O. Medalia, The molecular architecture of lamins in somatic cells. Nature 543, 261–264 (2017).

33. D. K. Shumaker, E. R. Kuczmarski, R. D. Goldman, The nucleoskeleton: lamins and actin are major players in essential nuclear functions. Curr Opin Cell Biol 15, 358–366 (2003).

34. X. Zheng, J. Hu, S. Yue, L. Kristiani, M. Kim, M. Sauria, J. Taylor, Y. Kim, Y. Zheng, Lamins Organize the Global Three-Dimensional Genome from the Nuclear Periphery. Mol Cell 71, 802–815 (2018).

35. T. Shimi, M. Kittisopikul, J. Tran, A. E. Goldman, S. A. Adam, Y. Zheng, K. Jaqaman, R. D. Goldman, Structural organization of nuclear lamins A, C, B1, and B2 revealed by superresolution microscopy. Mol Biol Cell 26, 4075–4086 (2015).

36. J. Nakayama, J. C. Rice, B. D. Strahl, C. D. Allis, S. I. Grewal, Role of histone H3 lysine 9 methylation in epigenetic control of heterochromatin assembly. Science 292, 110–113 (2001).

37. G. Liang, J. C. Lin, V. Wei, C. Yoo, J. C. Cheng, C. T. Nguyen, D. J. Weisenberger, G. Egger, D. Takai, F. A. Gonzales, P. A. Jones, Distinct localization of histone H3 acetylation and H3-K4 methylation to the transcription start sites in the human genome. Proc Natl Acad Sci U S A 101, 7357–7362 (2004).

38. L. Lau, Yin L. Lee, Steffen J. Sahl, T. Stearns, W. E. Moerner, STED Microscopy with Optimized Labeling Density Reveals 9-Fold Arrangement of a Centriole Protein. Biophysical Journal 102, 2926–2935 (2012).

39. X. Y. Shi, G. Garcia, J. C. Van De Weghe, R. McGorty, G. J. Pazour, D. Doherty, B. Huang, J. F. Reiter, Super-resolution microscopy reveals that disruption of ciliary transition-zone architecture causes Joubert syndrome. Nature Cell Biology 19, 1178-+ (2017).

40. A. J. Jin, R. Nossal, Rigidity of triskelion arms and clathrin nets. Biophys J 78, 1183–1194 (2000).

41. T. Kirchhausen, D. Owen, S. C. Harrison, Molecular structure, function, and dynamics of clathrin-mediated membrane traffic. Cold Spring Harb Perspect Biol 6, a016725 (2014).

42. G. Wen, M. Vanheusden, A. Acke, D. Valli, R. K. Neely, V. Leen, J. Hofkens, Evaluation of Direct Grafting Strategies via Trivalent Anchoring for Enabling Lipid Membrane and Cytoskeleton Staining in Expansion Microscopy. ACS Nano 14, 7860–7867 (2020).

43. B. Huang, S. A. Jones, B. Brandenburg, X. Zhuang, Whole-cell 3D STORM reveals interactions between cellular structures with nanometer-scale resolution. Nat Methods 5, 1047–1052 (2008).

44. M. Bates, B. Huang, X. Zhuang, Super-resolution microscopy by nanoscale localization of photo-switchable fluorescent probes. Curr Opin Chem Biol 12, 505–514 (2008).

45. E. D. Karagiannis, J. S. Kang, T. W. Shin, A. Emenari, S. Asano, L. Lin, E. K. Costa, A. H. Marblestone, N. Kasthuri, E. S. Boyden, Expansion Microscopy of Lipid Membranes. bioRxiv, doi: https://doi.org/10.1101/829903, (2019).

46. C. Y. Mao, M. Y. Lee, J. R. Jhan, A. R. Halpern, M. A. Woodworth, A. K. Glaser, T. J. Chozinski, L. Shin, J. W. Pippin, S. J. Shankland, J. T. Liu, J. C. Vaughan, Feature-rich covalent stains for super-resolution and cleared tissue fluorescence microscopy. Science Advances 6, EABA4542 (2020).

47. Magano, J. et al. Scalable and Cost-Effective Synthesis of a Linker for Bioconjugation with a peptide and a Monoclonal Antibody. Synthesis 46, 1399–1406 (2014).

48. Wosnick, J.H., Mello, C. M. & Swager, T. M. Synthesis and Application of Poly(phenylene Ethynylene)s for Bioconjugation: A Conjugated Polymer-Based Fluorogenic Probe for Proteases. J. Am. Chem. Soc. 127, 3400–3405 (2005).

49. McLaughlin, M., Mohareb, R. M. & Rapoport, H. An Efficient Procedure for the Preparation of 4-Substituted 5-Aminoimidazoles. J. Org. Chem. 68, 50–54 (2003).

50. Nainar, S. et al. Temporal Labeling of Nascent RNA Using Photoclick Chemistry in Live Cells. J. Am. Chem. Soc. 139, 8090–8093 (2017).

