## Supporting Information for "Label-retention expansion microscopy"

### Supplementary Materials

#### Supplementary Figures

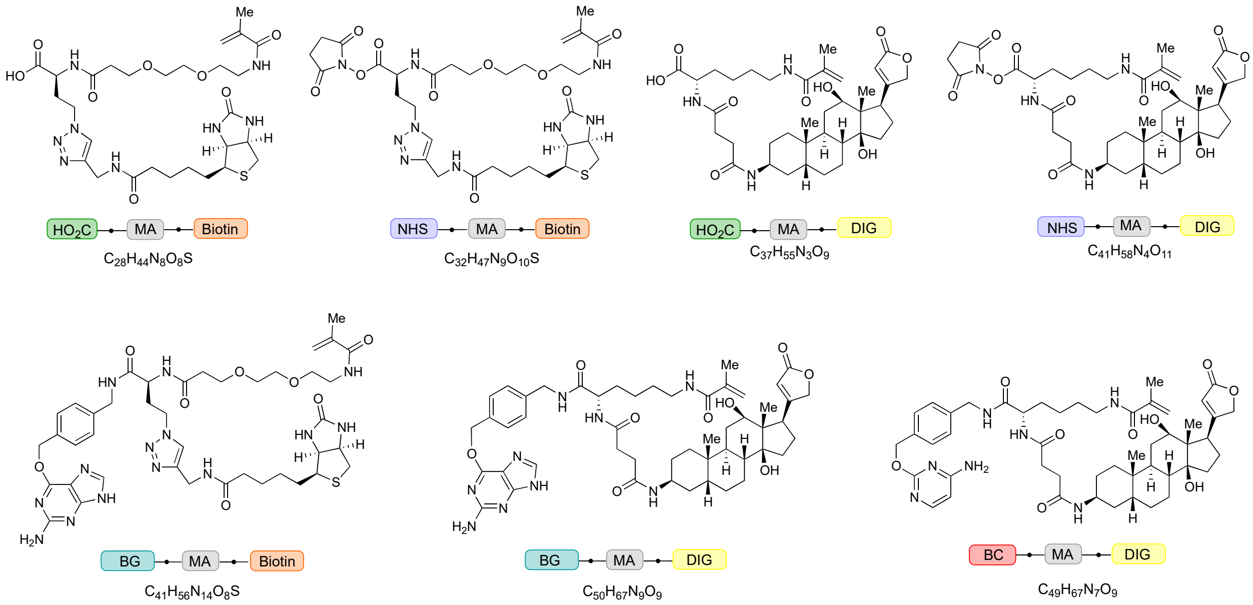


**Fig. S1.** Structures of trifunctional anchors HOOC/NHS-MA-Biotin, HOOC/NHS-MA-DIG, BG-MA-Biotin, BG-MA-DIG, and BC-MA-DIG.

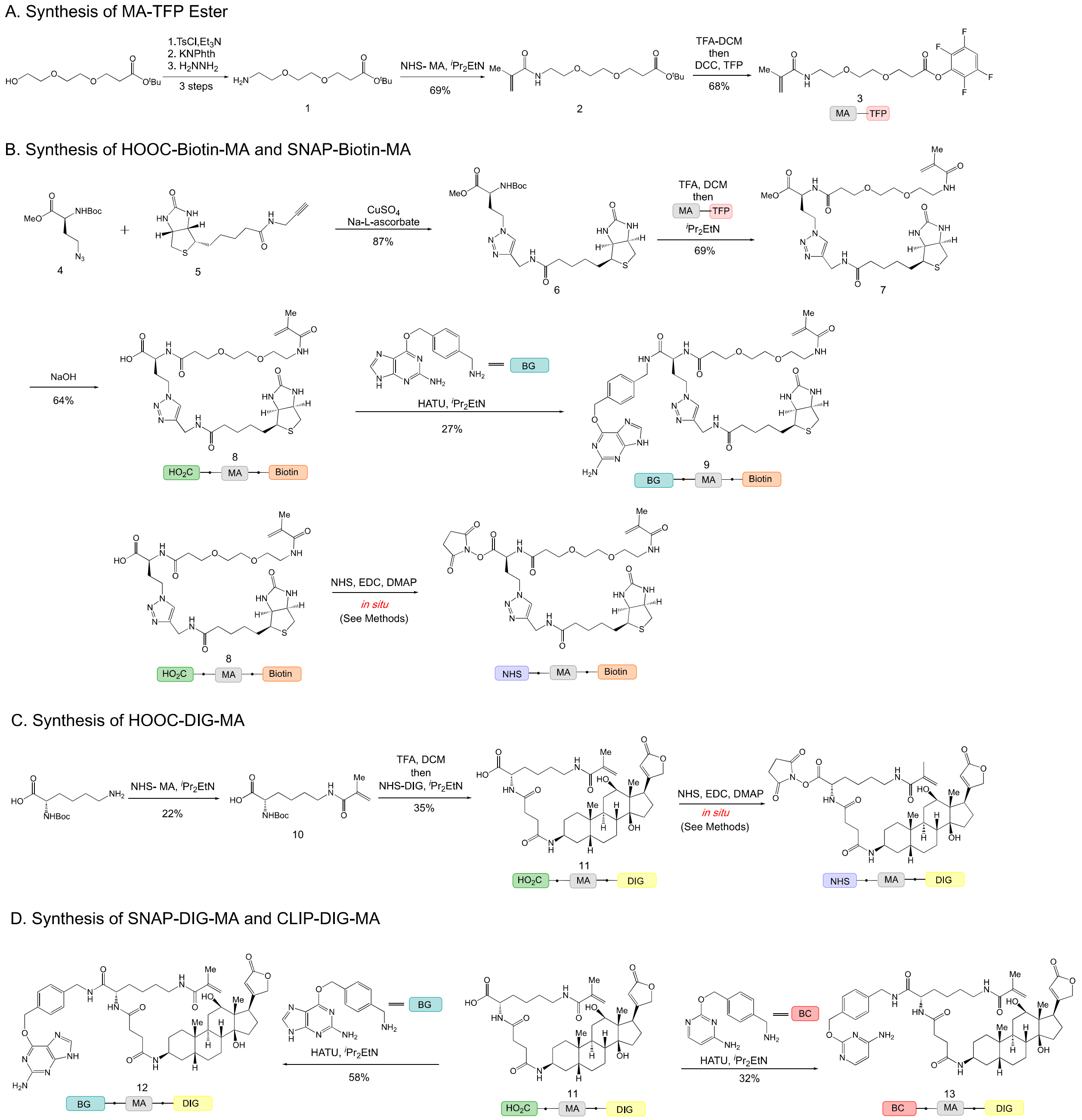


**Fig. S2.** Synthetic scheme of trifunctional anchors.

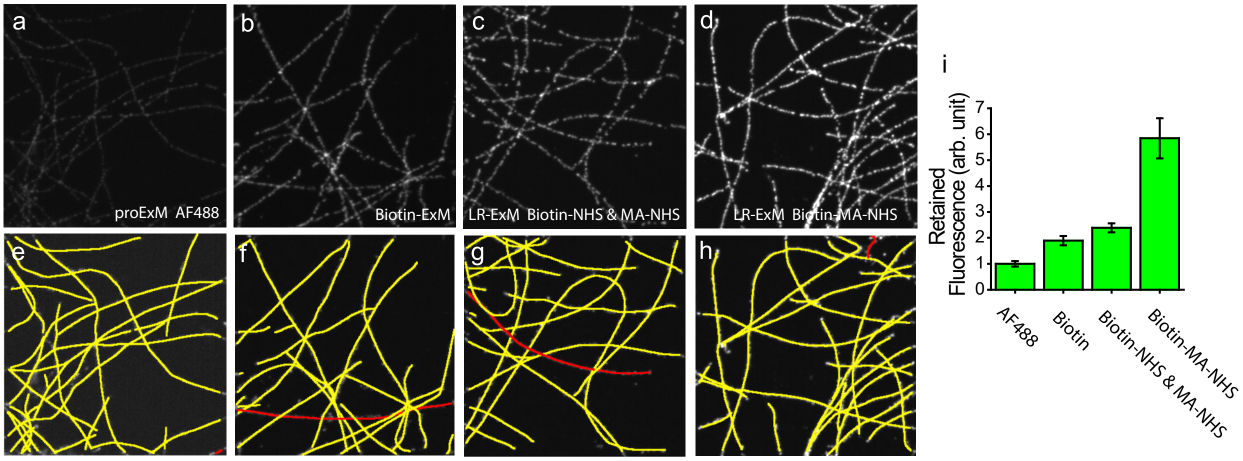


**Fig. S3.** Comparison of fluorescence intensities resulting from different ExM methods. Images of microtubules prepared with (a) proExM with Alexa Fluor 488 labeled secondary antibody, (b) biotin-ExM with the Biotin-NHS labeled secondary antibody, (c) LR-ExM with the Biotin-NHS and MA-NHS co-labeled secondary antibody, and (d) LR-ExM with the Biotin-MA-NHS labeled secondary antibody. Images a-d have the same contrast to show the relative brightness of the stain achieved in each case. Samples were processed side by side with the same immunostaining, digestion, and imaging conditions. The microtubules in images (a-d) are tracked and marked in yellow and red by a Fiji plugin JFilament as shown in images (e-h), respectively. (i) histogram of retained fluorescence in images (a-d). The retained fluorescence was normalized by the total length of microtubules in each image, and the ratios of AF488/Ab, Biotin/Ab, and AF488/Streptavidin. All samples were digested by incubating in 8 unit/mL proteinase K for 16 hours at room temperature.

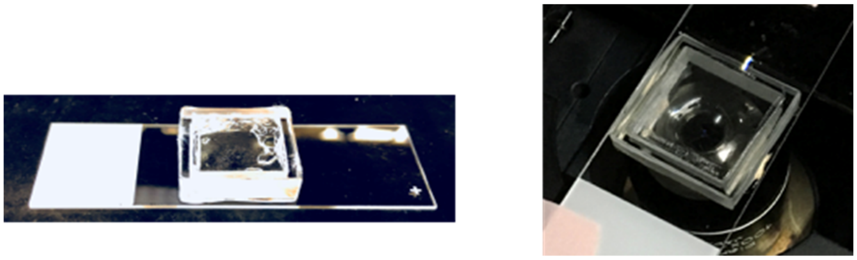

**Fig. S4.** 3D-printed chamber for drift reduction of hydrogel.

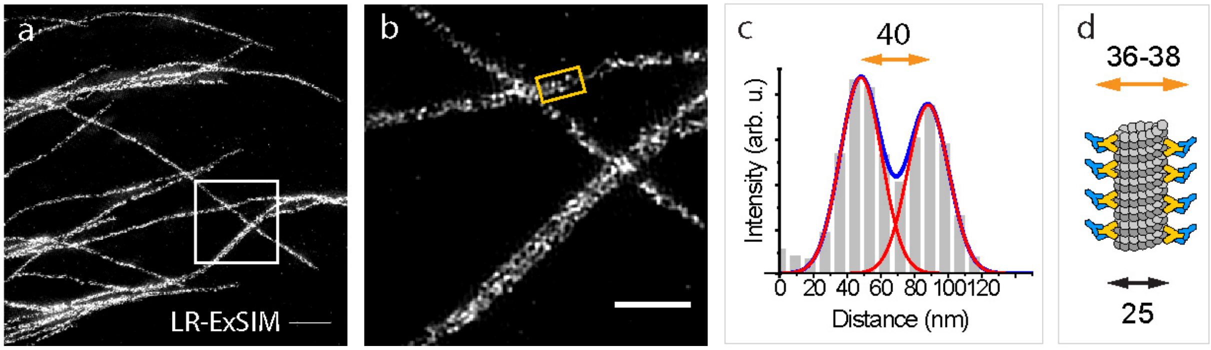


**Fig. S5.** LR-ExSIM of microtubules. (a) LR-ExSIM image of microtubules in a U2OS cell stained with antibody conjugated with NHS-MA-DIG anchors. (b) magnified image of (a). (c) the transverse profile of the microtubule in the gold box in image (b). (d) a schematic of the structure of immunostained microtubule. Scale bars: 1 μm (a) and 500 nm (b).

(c) The transverse profile of a microtubule in the final image showed two resolved peaks separated by about 40 nm, agreeing with the size of antibody-coated microtubules (d) previously measured by STORM[27,28]. By fitting the peaks to Gaussian functions, we calculated the resolution (FWHM) of LR-ExSIM to be 34 nm.

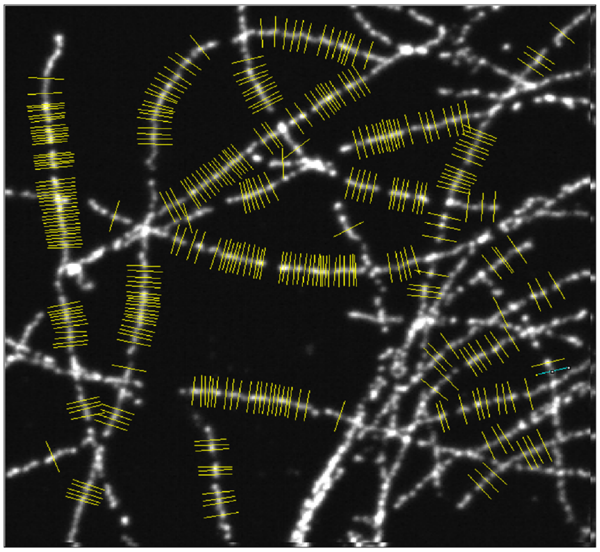

**Fig. S6.** Resolution measurement for LR-ExM confocal images. The transverse profiles of the microtubule cross sections marked in yellow were used to measure the resolution of LR-ExM using confocal microscope.

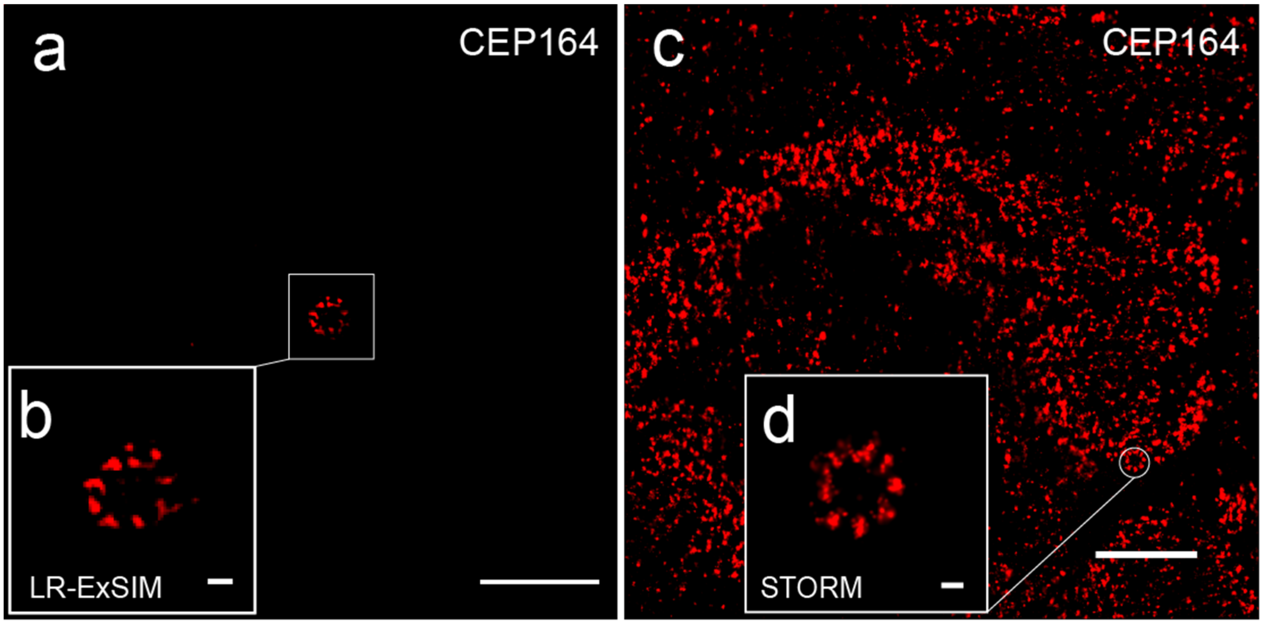


**Fig. S7. LR-ExSIM and STORM reveal structure of distal appendages with similar super resolution.** (a) LR-ExSIM image of Cep164 in distal appendages of a primary cilium of an expanded mouse embryonic fibroblast, indirectly immunostained with NHS-MA-Biotin secondary antibodies. The length expansion ratio is 4.2. (b) Magnified view of (a). (c) STORM image of Cep164 in distal appendages of motile cilia of a unexpanded multiciliated mouse tracheal epithelial cell. (d) Magnified view of (c). The same primary antibody was used for both images. Scale bars: 2 μm (a, c), and 100 nm (b, d).

#### Supplementary Materials and Methods

##### Reagents

Bifunctional linkers NHS-functionalized MA, biotin and digoxin were obtained from Lumiprobe (Hunt Valley, Maryland, USA). 1-Ethyl-3-(3-dimethylaminopropyl) carbodiimide (EDC), *N*-hydroxysuccinimide (NHS), 4-dimethylaminopyridine (DMAP), Trifluoroacetic acid (TFA), all the organic solvents were purchased from Sigma-Aldrich. HABA/Avidin reagent kit was obtained from Thermo Scientific. Nap™-5 sephadex G-25 size exclusion columns were ordered from GE Healthcare (GE Healthcare 17085301). Phosphate-buffered saline pH 7.4 (PBS) (catalog number: 10010-023) was ordered from Thermo Fisher Scientific (Waltham, MA, USA). Acetonitrile, dichlormethane, dimethylformamide, ether, and tetrahydrofuran to be used in anhydrous reaction mixtures were dried by passage through activated alumina columns immediately prior to use. Hexanes used were ≥85% *n*-hexane. Other commercial solvents and reagents were used as received, unless otherwise noted.

##### Trifunctional anchor synthesis and qualitative analysis

General: All reactions were performed in flame- or oven-dried glassware fitted with rubber septa under a positive pressure of nitrogen, unless otherwise noted. All reaction mixtures were stirred throughout the course of each procedure using Teflon-coated magnetic stir bars. Air- and moisture-sensitive liquids were transferred via syringe. Solutions were concentrated by rotary evaporation below 30 °C. Analytical thin-layer chromatography (TLC) was performed using glass plates pre-coated with silica gel (0.25-mm, 60-Å pore size, 230−400 mesh, SILICYCLE INC) impregnated with a fluorescent indicator (254 nm). TLC plates were visualized by exposure to ultraviolet light (UV) and then were stained by submersion in a basic aqueous solution of potassium permanganate or with an acidic ethanolic solution of anisaldehyde, followed by brief heating.

Materials: Dichloromethane (DCM), N,N-dimethylformamide (DMF), tetrahydrofuran (THF), ethyl ether, and acetonitrile to be used in anhydrous reaction mixtures were dried by passage through activated alumina columns immediately prior to use. Hexanes used were ≥85% n-hexane. Other commercial solvents and reagents were used as received, unless otherwise noted.

Instrumentation: Unless otherwise noted, proton nuclear magnetic resonance ($^1$H NMR) spectra and carbon nuclear magnetic resonance ($^{13}$C NMR) spectra were recorded on a 400 MHz Bruker Avance III HD 2-channel instrument NMR spectrometer at 23 °C. Proton chemical shifts are expressed in parts per million (ppm, δ scale) and are referenced to residual protium in the NMR solvent ($CHC1_3$: δ 7.26, DMSO-d5: δ 2.50, $CHD_2OD$: δ 3.31, HDO: δ 4.79). Carbon chemical shifts are expressed in parts per million (ppm, δ scale) and are referenced to the carbon resonance of the NMR solvent ($CDC1_3$: δ 77.16, DMSO-d6: δ 39.52, $CD_3OD$: δ 49.0). Data are represented as follows: chemical shift, multiplicity (s = singlet, d = doublet, t = triplet, q = quartet, dd = doublet of doublets, dt = doublet of triplets, sxt = sextet, m = multiplet, br = broad, app = apparent), integration, and coupling constant (J) in hertz (Hz). Reverse-phase preparative HPLC was carried out on a Waters Delta Prep 4000 preparative chromatography system [solvent A (0.1% TFA in Milli-Q® water) and B (0.1% TFA in acetonitrile), Gemini®-NX, 5μm, C18, 110Å, 30.00 mm i.d. x 100 mm), wavelength 220 nm with a gradient of 5–60% B over 17 min followed by 60-95% B over 2 min]. High-resolution mass spectra were obtained on a Waters Acquity UPLC/Xevo G2-XS QTOF mass spectrometer with ESI ionization (special thanks to Dr. Ziyang Zhang in the Shokat Laboratory for assistance). HPLC traces were obtained on an HP 1100 analytical HPLC [solvent A (0.1% TFA in Milli-Q® water) and B (0.1% TFA, 1% Milli-Q® water in acetonitrile), Jupiter®, 5μm, C4, 300Å, 4.6 mm i.d. x 250 mm), wavelength 220 nm with a gradient of 5–100% B over 45 min] (special thanks to Dr. Kui Zhang in the DeGrado Laboratory for assistance).

Amine 1[47]

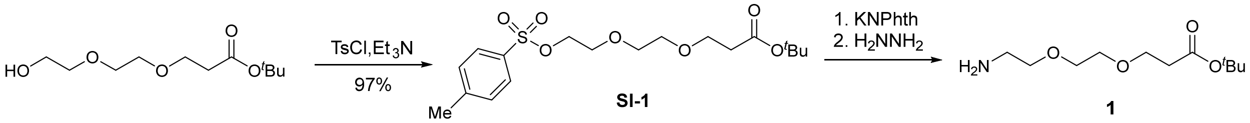


A 250-mL round-bottom flask containing tert-butyl 3-(2-(2hydroxyethoxy)ethoxy)propanoate (2.00 g, 8.54 mmol, 1 equiv) was evacuated and flushed with nitrogen (this process was repeated a total of 3 times). DCM (85.5 mL) was added followed by trimethylamine (4.7 mL, 34.1 mmol, 4.0 equiv), resulting in a colorless solution. The vessel was cooled to 0°C in an ice bath, and p-toluenesulfonyl chloride (4.07 g, 21.3 mmol, 2.5 equiv) was added in portion. The vessel was removed from the ice bath, and the system was allowed to warm to 23°C. After stirring overnight, the yellow solution was transferred to a separatory funnel and washed with water (2 x 100 mL) and brine (100 mL). The washed solution was dried ($Na_2SO_4$). The dried solution was filtered, and the filtrate was concentrated. The resulting crude residue was purified by flash chromatography (silica gel, eluent: EtOAc:hexanes = 1:3 to 1:1) to afford the tosylated product SI-1[48] (3.22 g, 97%) as a clear light yellow oil. The $^1$H NMR spectral data for tosylated product SI-1 is in agreement with the tabulated data published in reference 48.

DMF (82.4 mL) was added to a 250 mL round-bottom flask containing SI-1 (3.20 g, 8.24 mmol, 1 equiv), resulting in a light yellow solution. Potassium 1,3-dioxoisoindolin-2-ide (1.83 g, 9.88 mmol, 1.2 equiv) was added in one portion, resulting in a white suspension. The vessel was heated to 80°C with an oil bath. After 2 hours, the vessel was removed from the oil bath, and the system was allowed to cool to 23°C. The mixture was concentrated, and the residue was suspended between EtOAc (100 mL) and water (100 mL). The biphasic mixture was transferred to a separatory funnel, and the layers were separated. The organic layer was washed with water (100 mL) and brine (100 mL). The washed solution was dried ($Na_2SO_4$). The dried solution was filtered, and the filtrate was concentrated. The resulting crude residue was used in the next step without further purification.

Methanol was added to a 250 mL round-bottom flask containing the crude residue from the previous step (theoretical: 2.90 g, 7.98 mmol), resulting in a colorless solution. Hydrazine hydrate (0.774 mL, 16.0 mmol, 2 eq) was added. After 2 hours, the reaction vessel was equipped with a reflux condenser and heated to reflux with an oil bath. After another 2 hours, the vessel was removed from the oil bath, and the system was allowed to cool to 23°C. The mixture was concentrated, and DCM (100 mL) was added, resulting in a white suspension. The suspension was filtered through a pad of celite and the filter cake was washed with DCM (2 x 50 mL). The filtrate was concentrated, and the resulting amine 1 (1.59 g, 6.81 mmol) was used in the next step without further purification. The $^1$H NMR spectral data for amine 1 is in agreement with the tabulated data published in reference 47.

Methacrylamide 2

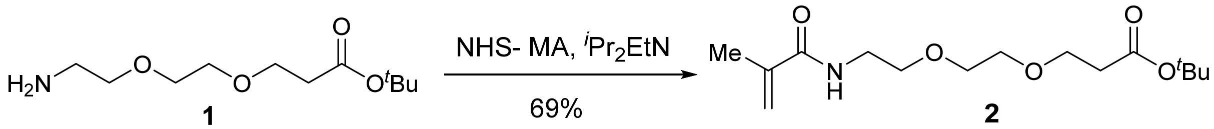


NHS-methacrylate (471 mg, 2.57 mmol, 1.2 equiv) was added to a 50-mL round-bottom flask containing amine 1 (500 mg, 2.14 mmol, 1 equiv). DCM (21 mL) was added. $^i$Pr$_2$EtN (0.75 mL, 4.29 mmol, 2.0 equiv) was then added dropwise. After 3 hours, the mixture was transferred to a separatory funnel and washed with water (2 x 25 mL) and brine (25 mL). The washed solution was dried ($Na_2SO_4$). The dried solution was filtered, and the filtrate was concentrated. The resulting crude residue was purified by flash chromatography (silica gel, eluent: EtOAc:hexanes = 1:1) to afford methylacrimide 2 (447 mg, 69%) as a colorless oil.

TLC (EtOAc:hexanes = 1:1): $R_f$ =0.26 (UV, anisaldehyde).

$^{1}H$ NMR (400 MHz, $CDCl_3$) δ 6.35 (s, 1H), 5.69 (t, *J* = 1.1 Hz, 1H), 5.31 (p, *J* = 1.5 Hz, 1H), 3.71 (t, *J* = 6.4 Hz, 2H), 3.66 – 3.54 (m, 6H), 3.55 – 3.46 (m, 2H), 2.49 (t, *J* = 6.5 Hz, 2H), 1.96 (s, 3H), 1.43 (s, 9H).
$^{13}C$ NMR (100 MHz, $CDCl_3$): δ 171.0, 168.5, 140.2, 119.6, 80.8, 70.4, 70.4, 69.9, 67.0, 39.5, 36.4, 28.2, 18.8.
HRMS-ESI m/z calcd for $C_{15}H_{27}NO_5$ $[M+Na]^+$ 324.1787, found 324.1801.

TFP ester 3

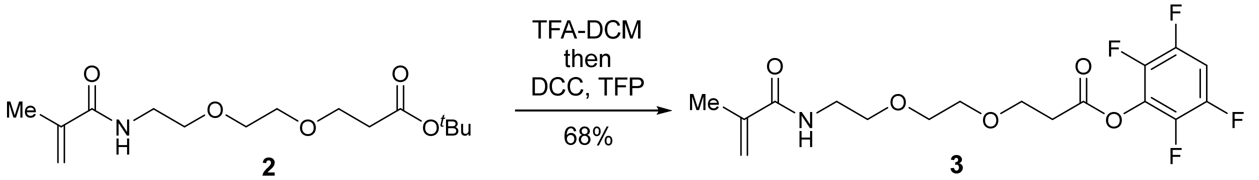


A 50-mL round-bottom flask containing methyacrylamide 2 (423 mg, 1.40 mmol, 1 equiv) was evacuated and flushed with nitrogen (this process was repeated a total of 3 times). DCM (21 mL) was added. Trifluoroacetic acid (7 mL) was then added dropwise. After stirring for 30 minutes, the mixture was concentrated. Toluene (10 mL) was added to the resulting crude residue, and the mixture was concentrated to remove residual trifluoroacetic acid (this process for repeated a total of 3 times). The residue was dissolved in DCM (14 mL), and 2,3,5,6-tetrafluorophenol (357 mg, 2.15 mmol, 1.5 equiv) and DCC (445 mg, 2.154 mmol, 1.5 equiv) were added sequentially. $^{i}Pr_2EtN$ (0.38 mL, 2.15 mmol, 1.5 equiv) was then added dropwise. After stirring overnight, the mixture was concentrated. The resulting crude was purified by flash chromatography (silica gel, eluent: EtOAc:hexanes = 1:1) to afford TFP-ester 3 (374 mg, 68%) as a colorless oil.
TLC (EtOAc:hexanes = 1:1): $R_f$ =0.23 (UV, anisaldehyde).
$^{1}H$ NMR (400 MHz, $CDCl_3$):δ 7.07 – 6.91 (m, 1H), 6.33 (s, 1H), 5.65 (s, 1H), 5.27 (s, 1H), 3.91 – 3.80 (m, 2H), 3.68 – 3.52 (m, 6H), 3.52 – 3.43 (m, 2H), 2.97 – 2.85 (m, 2H), 1.91 (s, 3H).

$^{13}C$ NMR (100 MHz, $CDCl_3$): δ 168.6 , 167.6 , 146.1 (dtd, *J* = 248.5, 11.8, 4.1 Hz), 140.6 (dddd, *J* = 250.6, 15.3, 4.8, 2.2 Hz), 140.1 , 129.5 (tt, *J* = 13.3, 3.9 Hz), 119.6 , 103.4 (t, *J* = 22.8 Hz), 70.6 , 70.2 , 69.8 , 66.1 , 39.4 , 34.5 , 18.6.
HRMS-ESI m/z calcd for $C_{17}H_{19}F_4NO_5$ $[M+H]^+$ 394.1279, found 344.1270.

Azide 4[49]

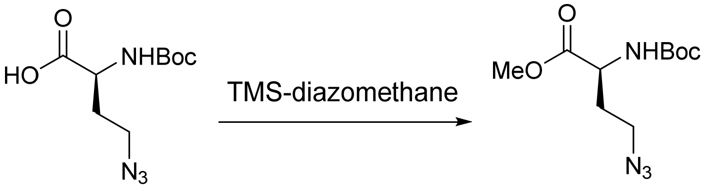


**4**

A 50-mL round-bottom flask containing (S)-4-azido-2-((tert-butoxycarbonyl)amino)butanoic acid (500 mg, 2.05 mmol, 1 equiv) was evacuated and flushed with nitrogen (this process was repeated a total of 3 times). A solution of MeOH and DCM (1:3, 22.8 mL) was added, resulting in a colorless solution. TMS-diazomethane (2 mL, 4.09 mmol, 2 equiv) was added dropwise. After 2 hours, the mixture was concentrated, and azide 4 (theoretical: 529 mg, 2.05 mmol) was used without further purification. The $^{1}H$ NMR spectral data for azide 4 is in agreement with the tabulated data published in reference 49.

Click product 6

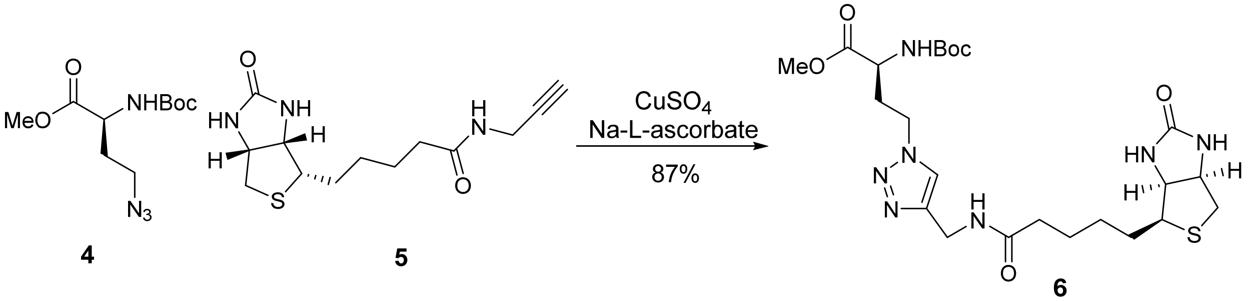


**4** **5** **6**

A 100-mL round-bottom flask containing azide 4 (529 mg, 2.05 mmol, 1.05 equiv) and alkyne 5[50] (549 mg, 1.95 mmol, 1 equiv) was evacuated and flushed with nitrogen (this process was repeated a total of 3 times). DCM (21 mL) was added, followed by adding a solution of copper (II) sulfate (31.1 mg, 195 μmol, 0.1 equiv) and sodium L-ascorbate (155 mg, 780 μmol, 0.4 equiv) in t-BuOH and $H_2O$ solvent (1:1, 19.5 mL). The resulting suspension was sonicated, resulting in a light yellow solution. After 2 hours, the solution was concentrated to reveal a yellow foam. The resulting crude residue was purified by flash chromatography (silica gel, eluent:MeOH:DCM = 1:9) to afford click product 6 (920 mg, 87%) as an off-white solid. TLC (MeOH:DCM = 1:9): $R_f$ =0.27 (anisaldehyde, $KMnO_4$).

$^{1}H$ NMR (400 MHz, DMSO-$d_6$): δ 8.28 (t, $J$ = 5.7 Hz, 1H), 7.86 (s, 1H), 7.44 (d, $J$ = 7.9 Hz, 1H), 6.41 (d, $J$ = 1.9 Hz, 1H), 6.36 (s, 1H), 4.51 – 4.21 (m, 5H), 4.16 – 4.04 (m, 1H), 4.00 – 3.89 (m, 1H), 3.61 (s, 3H), 3.09 (ddd, $J$ = 8.5, 6.2, 4.4 Hz, 1H), 2.82 (dd, $J$ = 12.4, 5.1 Hz, 1H), 2.57 (d, $J$ = 12.4 Hz, 1H), 2.31 – 2.16 (m, 1H), 2.14 – 2.00 (m, 3H), 1.70 – 1.11 (m, 15H).

$^{13}C$ NMR (100 MHz, DMSO-$d_6$): δ 172.3, 172.0, 162.7, 155.6, 145.1, 122.9, 78.6, 61.0, 59.2, 55.4, 52.0, 51.0, 46.2, 39.9, 35.0, 34.1, 31.3, 28.2, 28.2, 28.0, 25.2.

HRMS-ESI m/z calcd for $C_{23}H_{37}N_7O_6S$ $[M+H]^+$ 540.2606, found 540.2612.

Methyl ester 7

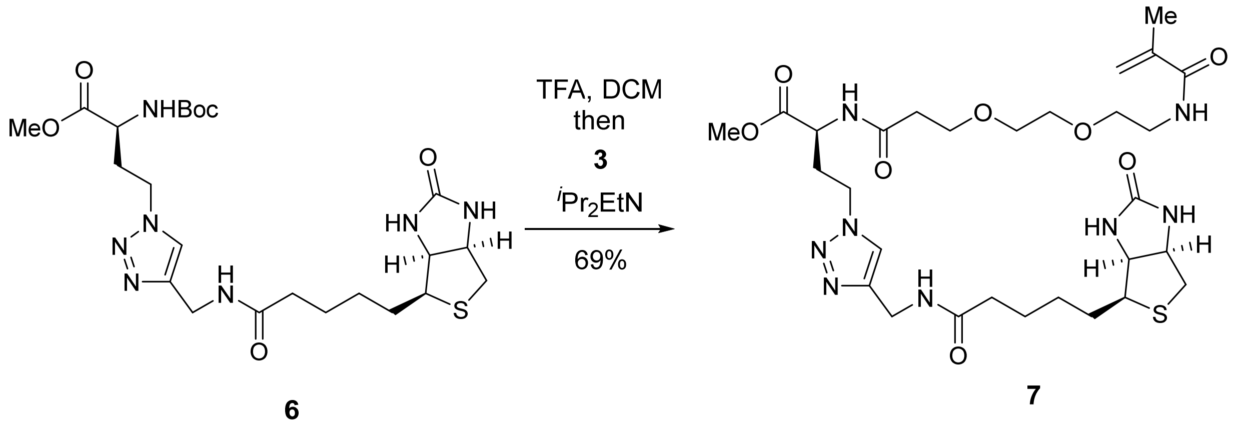

A 25-mL round-bottom flask containing click product 6 (200 mg, 371 μmol, 1 equiv) was evacuated and flushed with nitrogen (this process was repeated a total of 3 times). DCM (4 mL) was added followed by trifluoroacetic acid (4 mL). After 30 minutes, the mixture was concentrated. Toluene (10 mL) was added to the resulting crude residue, and the mixture was concentrated to remove residual trifluoroacetic acid (this process for repeated a total of 3 times). The residue was dissolved in DMF (8 mL), and a solution of TFP-ester 3 (160 mg, 408 μmol, 1.1 equiv) in DMF (4 mL) was added. $^{i}Pr_2EtN$ (0.32 mL, 1.85 mmol, 5 equiv) was then added dropwise. After 5 hours, the mixture was concentrated. The resulting crude residue was purified by flash chromatography (silica gel, eluent: MeOH:DCM = 1:19 to 1:9 with 1% ammonium hydroxide) to afford methyl ester 7 (170 mg, 69%) as a light pink oil.

TLC (MeOH:DCM = 1:9): $R_f$ =0.17 (UV, anisaldehyde).

$^1$H NMR (400 MHz, DMSO-$d_6$): δ 8.44 (d, $J$ = 7.8 Hz, 1H), 8.27 (t, $J$ = 5.7 Hz, 1H), 7.91 (t, $J$ = 5.7 Hz, 1H), 7.85 (s, 1H), 6.41 (s, 1H), 6.36 (s, 1H), 5.64 (s, 1H), 5.31 (t, $J$ = 1.6 Hz, 1H), 4.42 – 4.33 (m, 2H), 4.34 – 4.20 (m, 4H), 4.12 (ddd, $J$ = 7.8, 4.4, 1.9 Hz, 1H), 3.66 – 3.56 (m, 5H), 3.49 (s, 3H), 3.41 (t, $J$ = 6.1 Hz, 3H), 3.23 (q, $J$ = 6.0 Hz, 2H), 3.09 (ddd, $J$ = 8.5, 6.2, 4.4 Hz, 1H), 2.82 (dd, $J$ = 12.4, 5.0 Hz, 1H), 2.57 (d, $J$ = 12.4 Hz, 1H), 2.48 – 2.22 (m, 3H), 2.10 (q, $J$ = 7.2 Hz, 3H), 1.83 (s, 3H), 1.67 – 1.39 (m, 4H), 1.37 – 1.20 (m, 2H).

$^{13}$C NMR (100 MHz, DMSO-$d_6$): δ 172.0, 171.8, 170.6, 167.5, 162.7, 145.0, 139.8, 123.0, 119.1, , 69.5, 69.5, 68.8, 66.7, 61.0, 59.2, 55.4, 52.1, 49.3, 46.1, 39.9, 38.8, 35.9, 35.0, 34.1, 31.5, 28.2, 28.0, 25.2, 18.6.

HRMS-ESI m/z calcd for $C_{29}H_{46}N_8O_8S$ $[M+H]^+$ 667.3239, found 667.3251.

Carboxylic acid 8

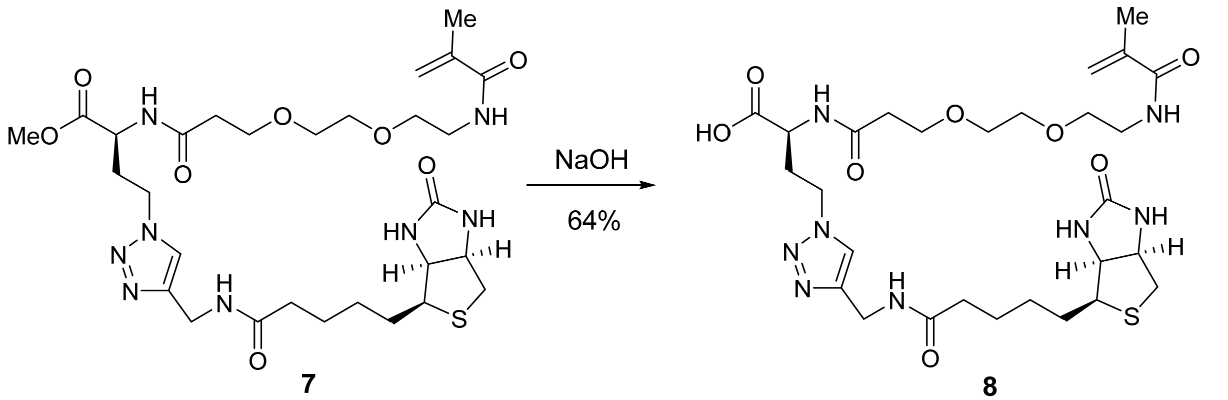


Water (5 mL) was added to a 20-mL scintillation vial containing methyl ester 7 (166 mg, 371 μmol, 1 equiv). Sodium hydroxide (49.8 mg, 1.24 mmol, 5 equiv) was added. After 1 hour, the reaction was quenched with 1N HCl (1.6 mL) and concentrated. The resulting crude residue was purified by reverse-phase high-performance liquid chromatography to afford carboxylic acid 8 (103 mg, 64%) as a white powder.

TLC (MeOH:DCM = 1:1): $R_f$ =0.17 (UV, anisaldehyde).

$^{1}$H NMR (400 MHz, $D_2O$): δ 7.90 (s, 1H), 5.67 (s, 1H), 5.43 (s, 1H), 4.63 – 4.56 (m, 1H), 4.57 – 4.48 (m, 2H), 4.46 (s, 2H), 4.38 (dd, *J* = 7.9, 4.5 Hz, 1H), 4.31 (dd, *J* = 9.7, 4.6 Hz, 1H), 3.79 (t, *J* = 6.1 Hz, 2H), 3.68 (s, 4H), 3.63 (t, *J* = 5.4 Hz, 2H), 3.43 (t, *J* = 5.4 Hz, 2H), 3.28 (p, *J* = 5.6 Hz, 1H), 2.98 (dd, *J* = 13.0, 5.0 Hz, 1H), 2.77 (d, *J* = 13.0 Hz, 1H), 2.63 – 2.50 (m, 3H), 2.40 – 2.25 (m, 3H), 1.91 (s, 3H), 1.77 – 1.43 (m, 4H), 1.43 – 1.24 (m, 2H).

$^{13}$C NMR (100 MHz, $D_2O$): δ 176.7, 174.4, 174.0, 171.9, 165.3, 144.7, 139.0, 124.1, 121.0, 69.6, 69.4, 68.8, 66.6, 62.0, 60.2, 55.4, 40.0, 47.0, 39.7, 39.1, 35.7, 35.2, 34.2, 30.8, 27.7, 27.6, 25.0, 17.7.

HRMS-ESI m/z calcd for $C_{28}H_{44}N_8O_8S$ $[M+H]^+$ 653.3083, found 653.3079.

Trifunctional anchor 9

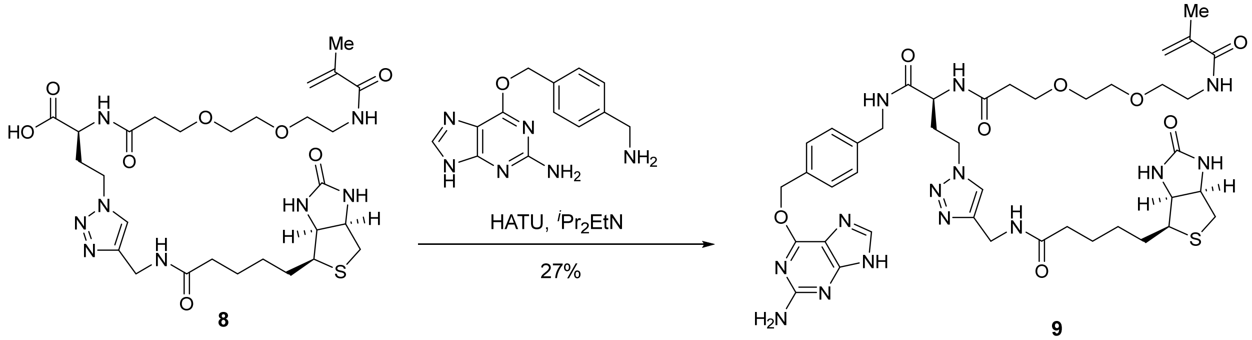


A 10-mL round-bottom flask containing carboxylic acid 8 (20 mg, 30.6 μmol, 1 equiv) and 6-((4-

(aminomethyl)benzyl)oxy)-9H-purin-2-amine (12.4 mg, 46.0 µmol, 1.5 equiv) was evacuated and flushed with nitrogen (this process was repeated a total of 3 times). DMF (2 mL) was added. $^{i}Pr_2EtN$ (13 µL, 76.6 µmol, 2.5 equiv) was then added dropwise. After the solid reactants dissolved, HATU (23.3 mg, 61.3µmol, 2 equiv) was added, resulting in a yellow solution. After 2 hours, the mixture was concentrated to reveal a yellow oil. The resulting crude residue was purified by reverse-phase high-performance liquid chromatography to afford trifunctional anchor 9 (7.5 mg, 27%) as a white powder.

TLC (MeOH:DCM = 1:4): $R_f$ =0.18 (UV, anisaldehyde).

$^{1}H$ NMR (400 MHz, MeOD): δ 8.38 (s, 1H), 7.83 (s, 1H), 7.50 (d, *J* = 8.1 Hz, 2H), 7.32 (d, *J* = 8.1 Hz, 2H), 5.68 (s, 1H), 5.64 (s, 2H), 5.38 – 5.34 (m, 1H), 4.52 – 4.32 (m, 8H), 4.27 (dd, *J* = 7.9, 4.5 Hz, 1H), 3.84 – 3.68 (m, 2H), 3.60 – 3.53 (m, 4H), 3.50 (t, *J* = 5.8 Hz, 2H), 3.35 (t, *J* = 5.8 Hz, 2H), 3.21 – 3.13 (m, 1H), 2.90 (ddd, *J* = 12.8, 5.0, 2.7 Hz, 1H), 2.69 (dd, *J* = 12.8, 3.0 Hz, 1H), 2.61 – 2.37 (m, 3H), 2.31 – 2.15 (m, 3H), 1.91 (s, 3H), 1.80 – 1.51 (m, 4H), 1.47 – 1.33 (m, 2H).

$^{13}C$ NMR (100 MHz, MeOD): δ 176.0, 174.3, 171.3, 166.1, 161.1, 153.4, 146.3, 143.9, 141.2, 140.7, 135.3, 130.3, 128.7, 124.7, 120.6, 107.7, 71.3, 71.2, 71.0, 70.4, 68.3, 63.3, 61.6, 57.0, 52.2, 49.7, 49.5, 49.3, 49.1, 48.1, 43.8, 41.0, 40.4, 37.6, 36.5, 35.6, 33.4, 29.7, 29.4, 26.7, 18.8.

HRMS-ESI m/z calcd for $C_{41}H_{56}N_{14}O_8S$ $[M+H]^+$ 905.4206, found 905.4189.

Methyacrylamide 10

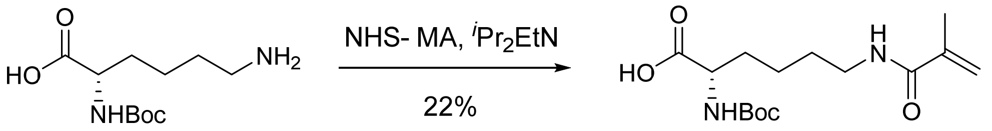


**10**

A 10-mL round-bottom flask containing (tert-butoxycarbonyl)-L-lysine (200 mg, 812 µmol, 1 equiv) and NHS-methacrylate (178 mg, 974 µmol, 1.2 equiv) was evacuated and flushed with nitrogen (this process was repeated a total of 3 times). DMF (8 mL) was added. $^{i}Pr_2EtN$ (0.28 mL, 1.62 mmol, 2 equiv) was then added dropwise, resulting in a white suspension. After stirring overnight, the mixture was concentrated to reveal a yellow oil. The resulting crude residue was purified by flash chromatography (silica gel, eluent: MeOH:DCM=1:19), to yield an oil contaminated with n-hydroxysuccinamide. The oil was dissolved in DCM (10 mL), and then washed with water (3 x 10 mL) and brine (10 mL). The washed solution was dried ($Na_2SO_4$). The dried solution was filtered, and the filtrate was concentrated to yield methacrylamide 10 (71.4 mg, 22%) as a clear oil.

TLC (MeOH:DCM = 1:9): $R_f$ =0.19 (UV, anisaldehyde).

$^{1}H$ NMR (400 MHz, $CDCl_3$): δ 8.70 (brs, 1H), 6.31 (brs, 1H), 5.69 (s, 1H), 5.38 (brs, 1H), 5.31 (s, 1H), 4.23 (brs, 1H), 3.38-3.18 (m, 2H), 1.93 (s, 3H), 1.84 (bs, 1H), 1.76 – 1.49 (m, 4H), 1.42 (s, 9H), 1.30 – 1.19 (m, 1H).
$^{13}C$ NMR (100 MHz, $CDCl_3$): δ 176.0, 169.2, 156.0, 139.9, 120.1, 80.1, 53.6, 39.5, 32.2, 29.0, 28.5, 22.6, 18.8.
HRMS-ESI m/z calcd for $C_{15}H_{26}N_2O_5$ $[M+Na]^+$ 337.1739, found 337.1774.

Carboxylic Acid 11

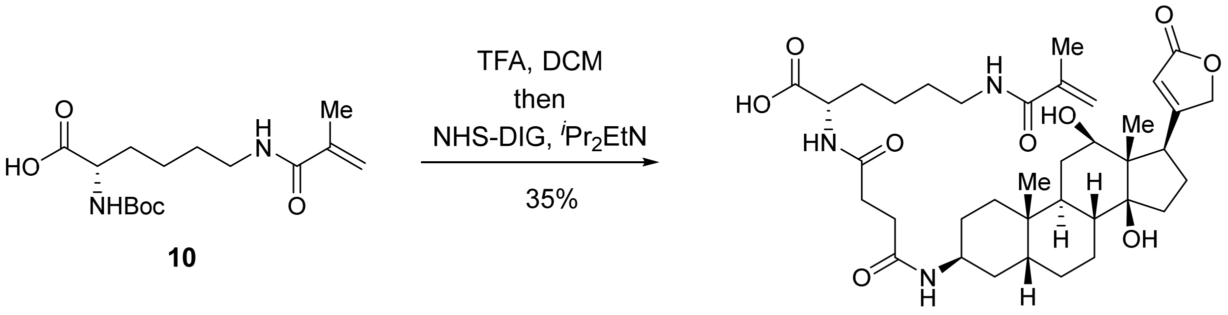


**10** **11**

A 5-mL round-bottom flask containing methacrylamide 10 (2.6 mg, 8.3 μmol, 1 equiv) was evacuated and flushed with nitrogen (this process was repeated a total of 3 times). DCM (1 mL) was added. Trifluoroacetic acid (1 mL) was then added dropwise. After 30 minutes, the mixture was concentrated. Toluene was added to the resulting crude residue, and the mixture was concentrated to remove residual trifluoroacetic acid (this process for repeated a total of 3 times). The residue was dissolved in DCM, and 3-amino-3-deoxydigoxigenin hemisuccinamide, succinimidyl ester (5.0 mg, 8.5 μmol, 1.0 equiv) was added in one portion. $^{i}Pr_2EtN$ (7.2 μL, 41 μmol, 5.0 equiv) was then added dropwise. After stirring overnight, the mixture was concentrated. The resulting crude residue was purified by reverse-phase high-performance liquid chromatography to afford carboxylic acid 11 (2.0 mg, 35%) as a white powder.
TLC (MeOH:DCM = 1:4): $R_f$ =0.12 (UV, anisaldehyde).
$^{1}H$ NMR (400 MHz, MeOD): δ 5.91 (brt, $J$ = 1.8 Hz, 1H), 5.67 (brt, $J$ = 1.0 Hz, 1H), 5.37 – 5.34 (m, 1H), 5.03 – 4.87 (m, 2H), 4.37 (dd, $J$ = 8.9, 4.9 Hz, 1H), 4.07 (bs, 1H), 3.44 – 3.32 (m, 2H), 3.24 (t, $J$ = 6.9 Hz, 2H), 2.67 – 2.36 (m, 4H), 2.22 – 1.07 (m, 25H), 1.93 (s, 3H), 1.00 (s, 3H), 0.79 (s, 3H).
HRMS-ESI m/z calcd for $C_{37}H_{55}N_3O_9$ $[M+H]^+$ 686.4018, found 686.4000.

Trifunctional anchor 12

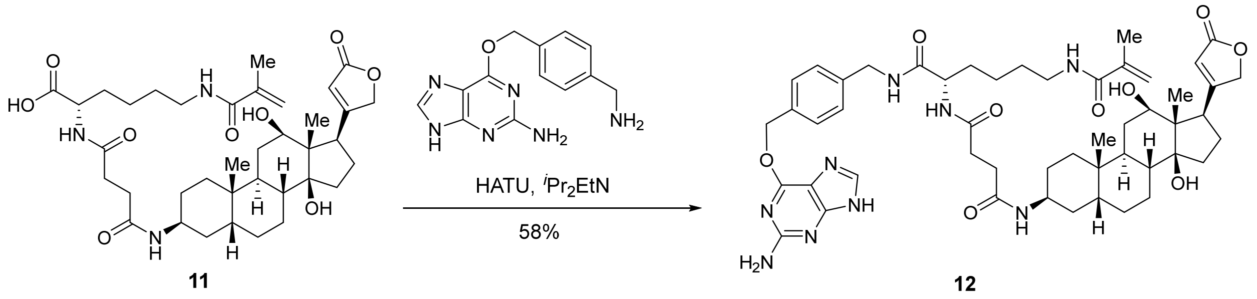


A 5-mL round-bottom flask containing carboxylic acid 11 (1.5 mg, 2.2 μmol, 1 equiv) was evacuated and flushed with nitrogen (this process was repeated a total of 3 times). DMF (0.3 mL) was added. $^{i}Pr_2EtN$ (0.95 μL, 5.5 μmol, 2.5 equiv) was then added dropwise. After stirring overnight, 6-((4-(aminomethyl)benzyl)oxy)-7H-purin-2-amine (0.89 mg, 3.3 μmol, 1.5 equiv) and HATU(1.7 mg, 4.4 μmol, 2 equiv) were added in succession. After 2 hours, the mixture was concentrated. The resulting crude was purified by reverse-phase high-performance liquid chromatography to afford trifunctional anchor 12 (1.2 mg, 58%) as a white powder.

TLC (MeOH:DCM = 1:4): $R_f$ =0.53 (UV, anisaldehyde).

$^{1}$H NMR (400 MHz, MeOD) δ 8.66 (t, *J* = 6.0 Hz, 1H), 8.28 (s, 1H), 7.49 (d, *J* = 8.0 Hz, 2H), 7.32 (d, *J* = 8.0 Hz, 2H), 5.90 (s, 1H), 5.67 (s, 1H), 5.62 (s, 2H), 5.37 – 5.33 (m, 1H), 5.03 – 4.87 (m, 2H), 4.46 – 4.34 (m, 2H), 4.30 (dd, *J* = 9.3, 4.7 Hz, 1H), 3.97 (bs, 1H), 3.44 – 3.32 (m, 2H), 3.23 (t, *J* = 7.0 Hz, 2H), 2.92 – 2.85 (m, 1H), 2.71 – 2.35 (m, 5H), 2.24 – 1.06 (m, 25H), 1.93 (s, 3H), 0.97 (s, 3H), 0.78 (s, 3H).

HRMS-ESI m/z calcd for $C_{50}H_{67}N_9O_9$ $[M+H]^+$ 938.5142, found 938.5134.

Trifunctional anchor 13

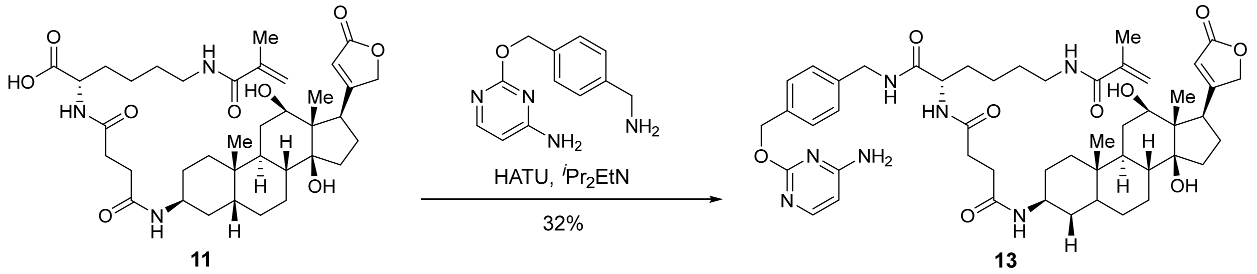


A 5-mL round-bottom flask containing carboxylic acid 11 (1.5 mg, 2.2 μmol, 1 equiv) was evacuated and flushed with nitrogen (this process was repeated a total of 3 times). DMF (0.3 mL) was added. $^{i}Pr_2EtN$ (0.95 μL, 5.5 μmol, 2.5 equiv) was then added dropwise. After stirring overnight, 2-((4-(aminomethyl)benzyl)oxy)pyrimidin-4-amine (0.76 mg, 3.3 μmol, 1.5 equiv) and HATU(1.7 mg, 4.4 μmol, 2 equiv) were added in succession. After 2 hours, the mixture was concentrated. The resulting crude was purified by reverse-phase high-performance liquid chromatography to afford trifunctional anchor 13 (0.6 mg, 32%) as a white powder.

TLC (MeOH:DCM = 1:4): $R_f$ =0.53 (UV, anisaldehyde).

$^{1}$H NMR (400 MHz, MeOD) δ 8.67 (t, *J* = 6.0 Hz, 1H), 7.87 (d, *J* = 7.1 Hz, 1H), 7.45 (d, *J* = 8.0 Hz, 2H), 7.33 (d, *J* = 8.0 Hz, 2H), 6.39 (d, *J* = 7.1 Hz, 1H), 5.91 (s, 1H), 5.67 (s, 1H), 5.51 (s, 2H), 5.37 – 5.33 (m, 1H), 5.03 – 4.87 (m, 2H), 4.45 – 4.37 (m, 2H), 4.30 (dd, *J* = 9.3, 4.6 Hz, 1H), 3.99 (bs, 1H), 3.44 – 3.32 (m, 2H), 3.23 (t, *J* = 7.0 Hz, 2H), 2.89 (d, *J* = 11.0 Hz, 1H), 2.69 – 2.39 (m, 5H), 2.20 – 1.15 (m, 25H), 1.93 (s, 3H), 0.98 (s, 3H), 0.78 (s, 3H).

HRMS-ESI m/z calcd for $C_{49}H_{67}N_7O_9$ $[M+H]^+$ 898.5008, found 898.5031.

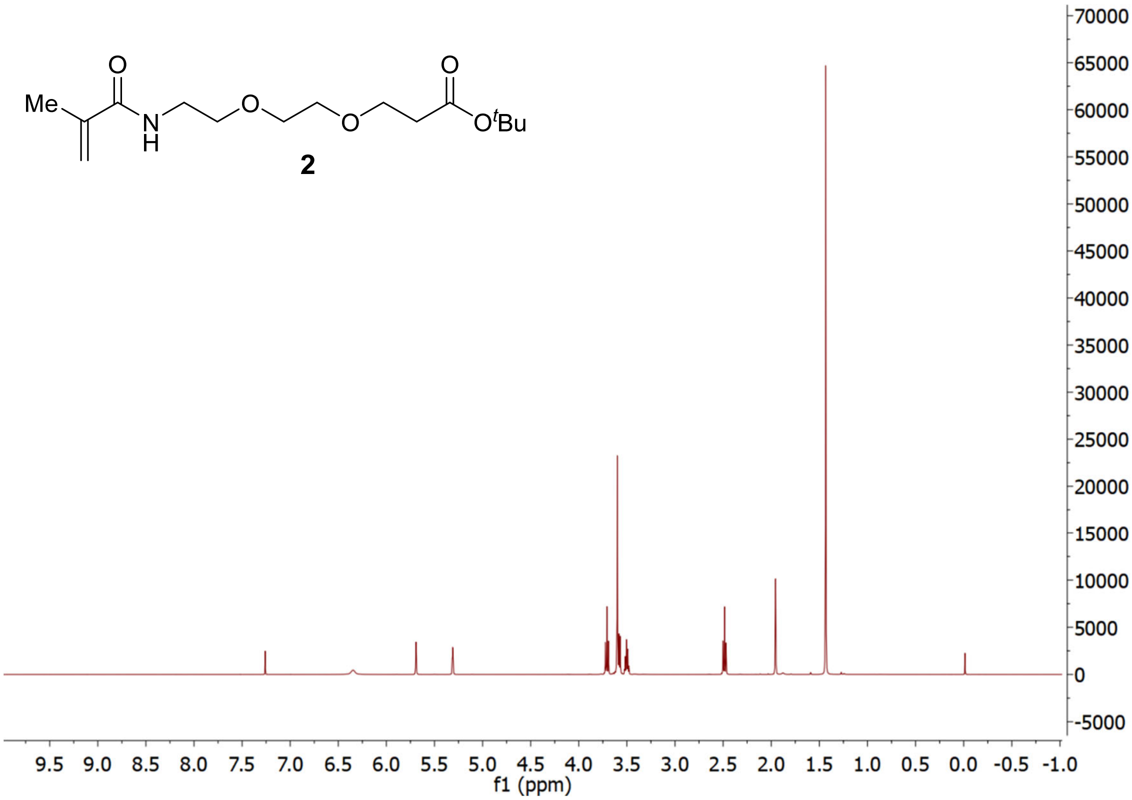

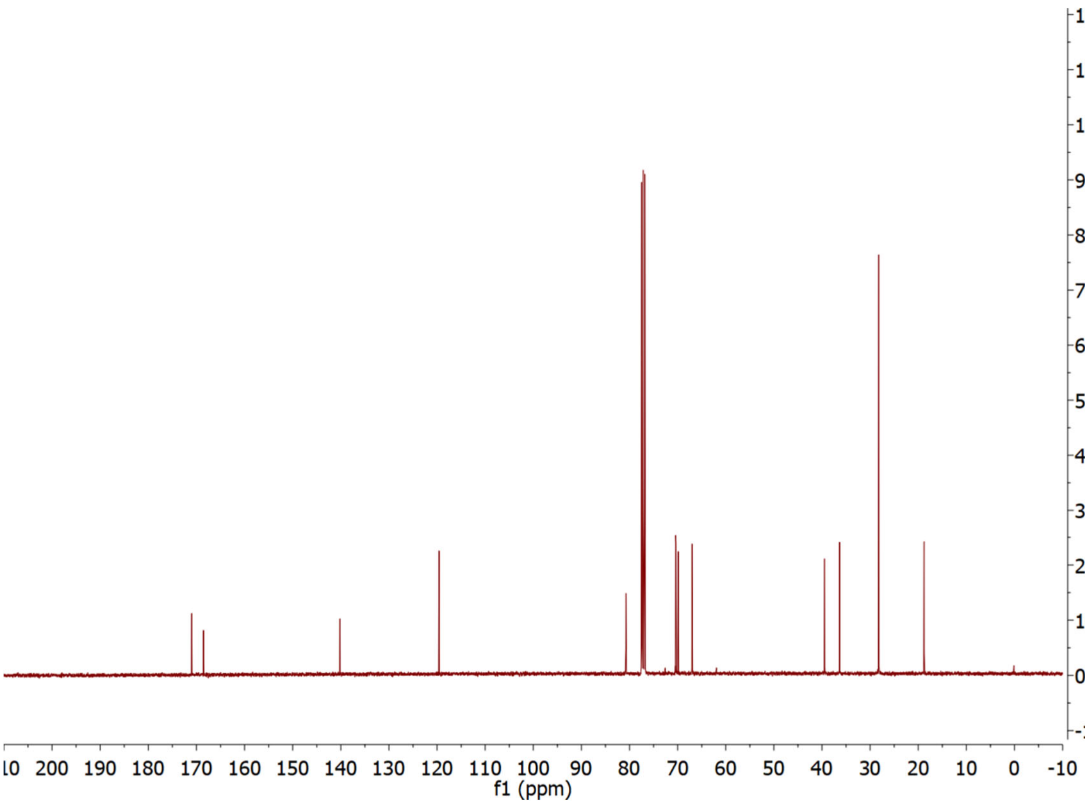

12000
11000
10000
9000
8000
7000
6000
5000
4000
3000
2000
1000
0
-1000
10 200 190 180 170 160 150 140 130 120 110 100 90 80 70 60 50 40 30 20 10 0 -10
f1 (ppm)


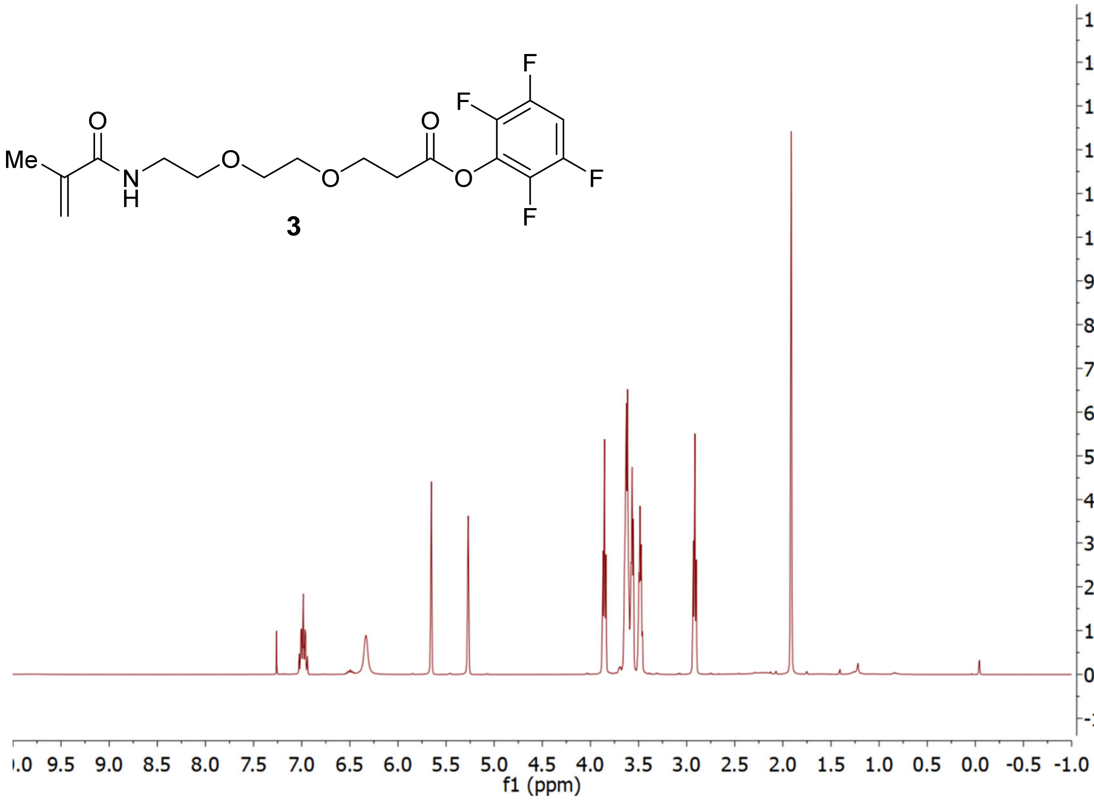

Me
O
N
H
O
O
O
O
F
F
F
F
3
15000
14000
13000
12000
11000
10000
9000
8000
7000
6000
5000
4000
3000
2000
1000
0
-1000
.0 9.5 9.0 8.5 8.0 7.5 7.0 6.5 6.0 5.5 5.0 4.5 4.0 3.5 3.0 2.5 2.0 1.5 1.0 0.5 0.0 -0.5 -1.0
f1 (ppm)

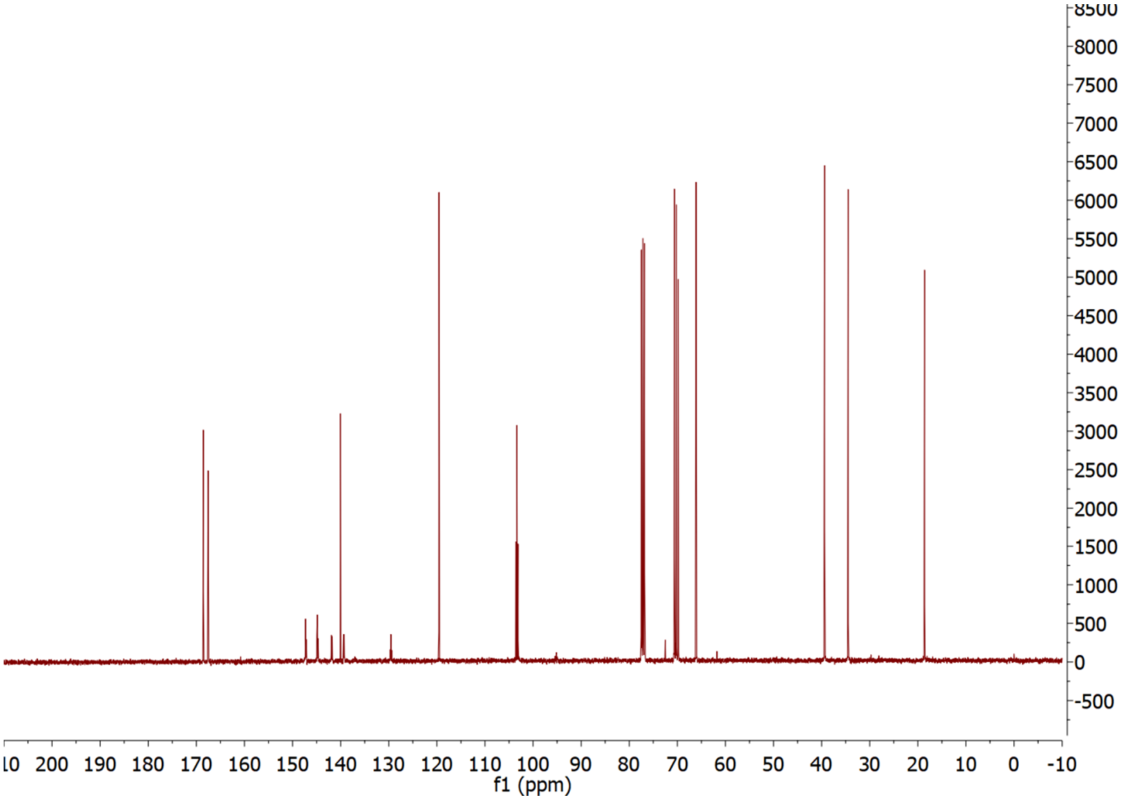


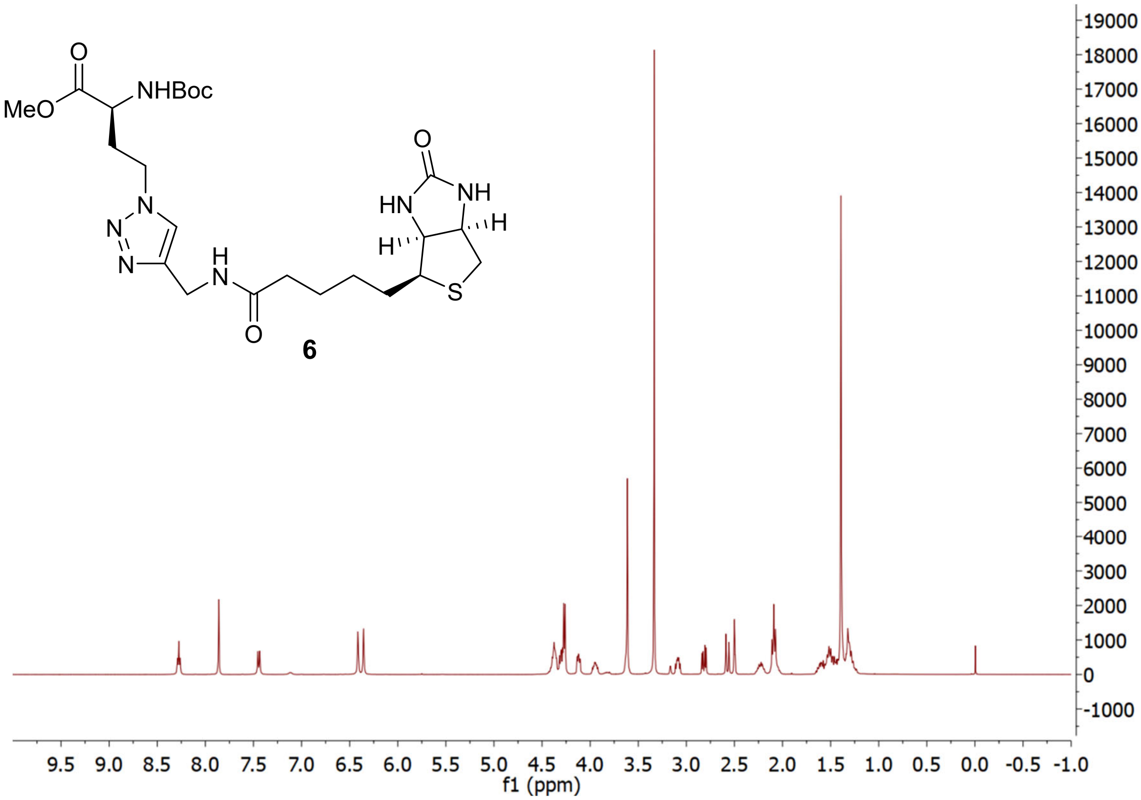


6

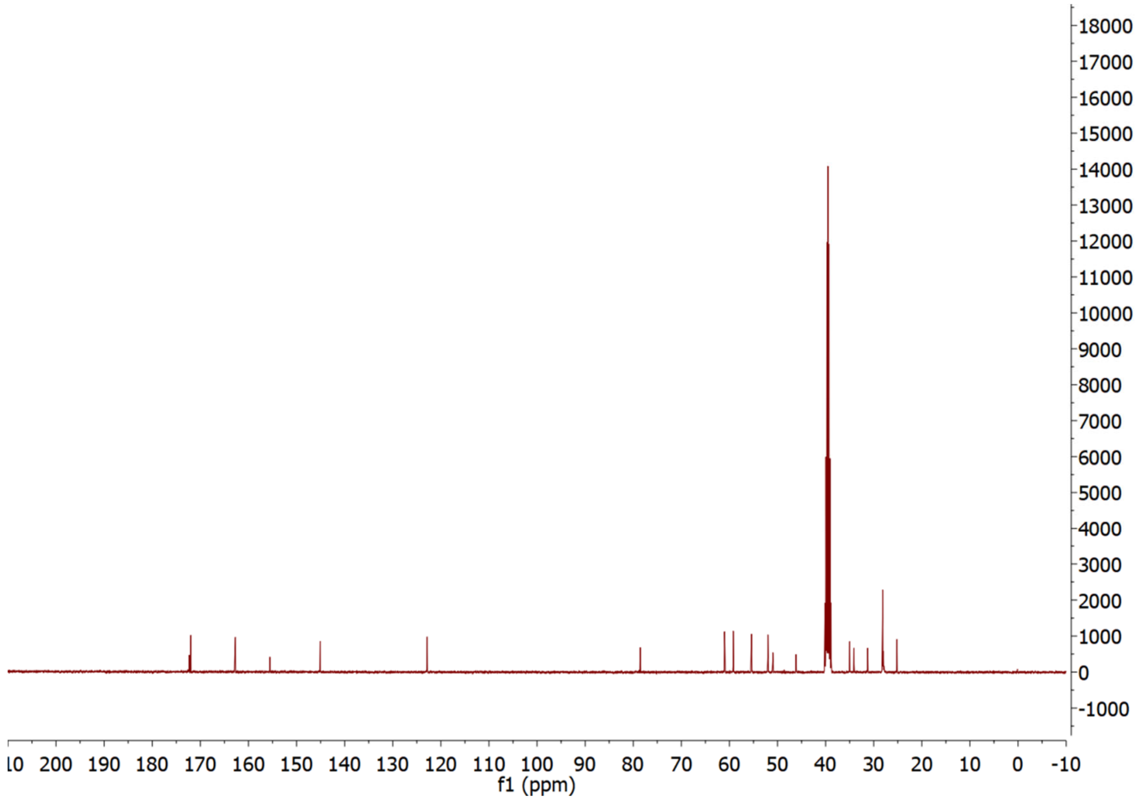

18000 17000 16000 15000 14000 13000 12000 11000 10000 9000 8000 7000 6000 5000 4000 3000 2000 1000 0 -1000
10 200 190 180 170 160 150 140 130 120 110 100 90 80 70 60 50 40 30 20 10 0 -10
f1 (ppm)


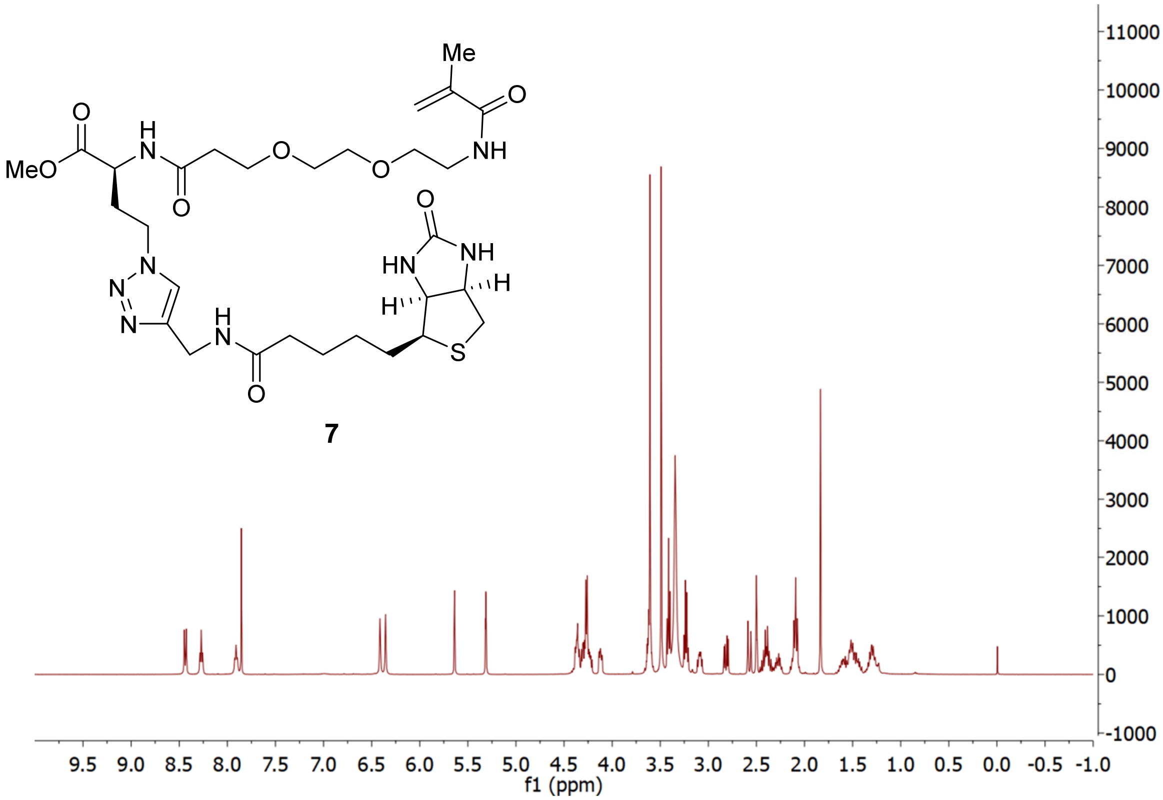

Me
O
O
H
N
MeO
O
O
NH
O
N
N
N
N
H
N
O
O
HN
NH
H
H
S
11000 10000 9000 8000 7000 6000 5000 4000 3000 2000 1000 0 -1000
9.5 9.0 8.5 8.0 7.5 7.0 6.5 6.0 5.5 5.0 4.5 4.0 3.5 3.0 2.5 2.0 1.5 1.0 0.5 0.0 -0.5 -1.0
f1 (ppm)


**7**

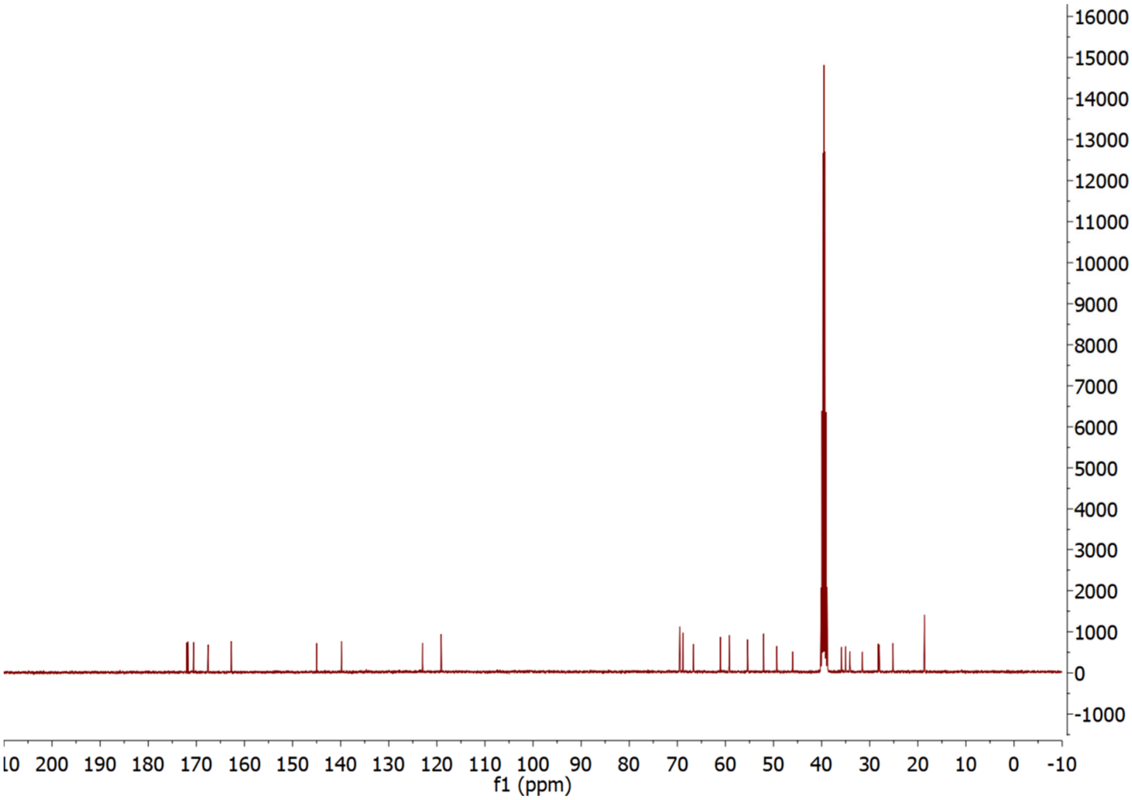


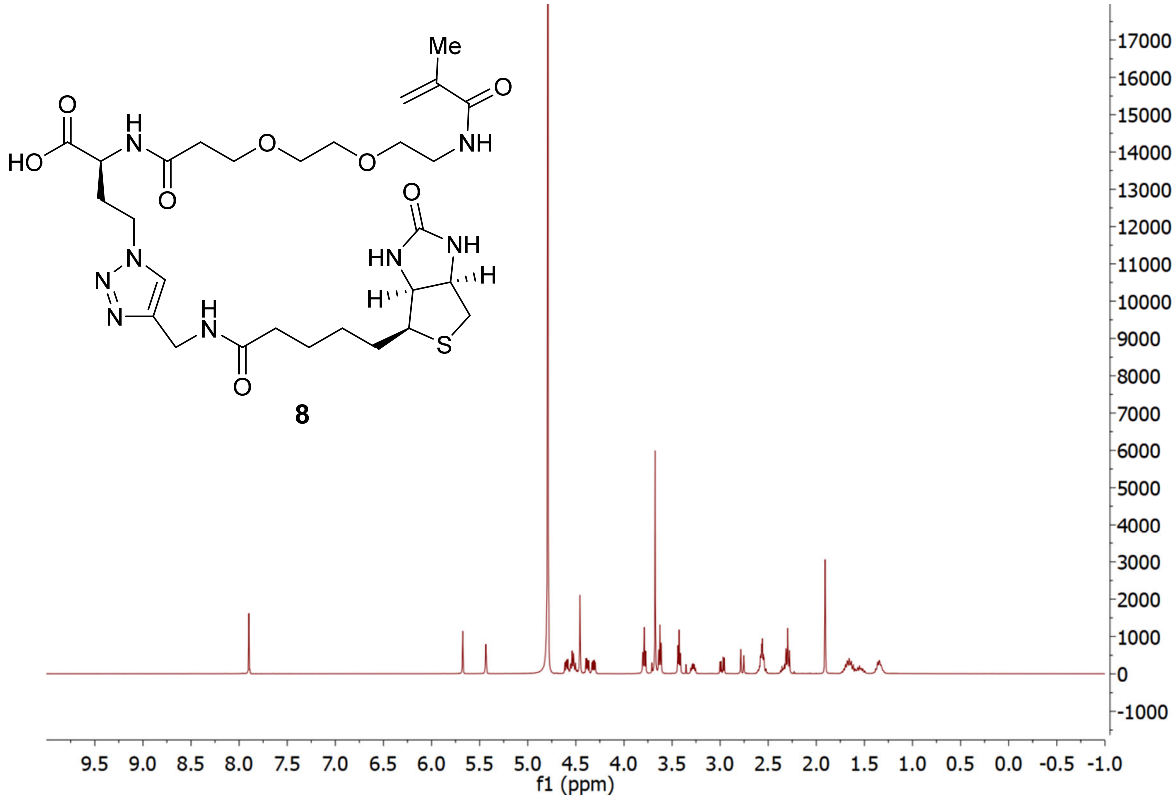


**8**

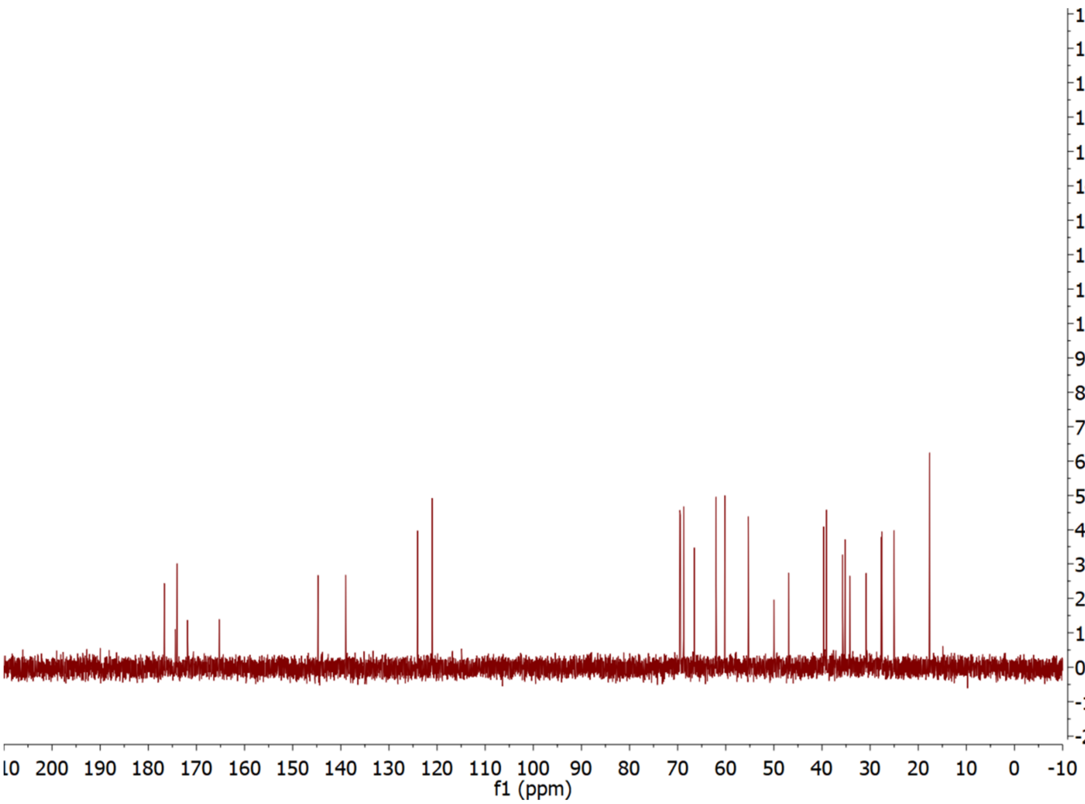


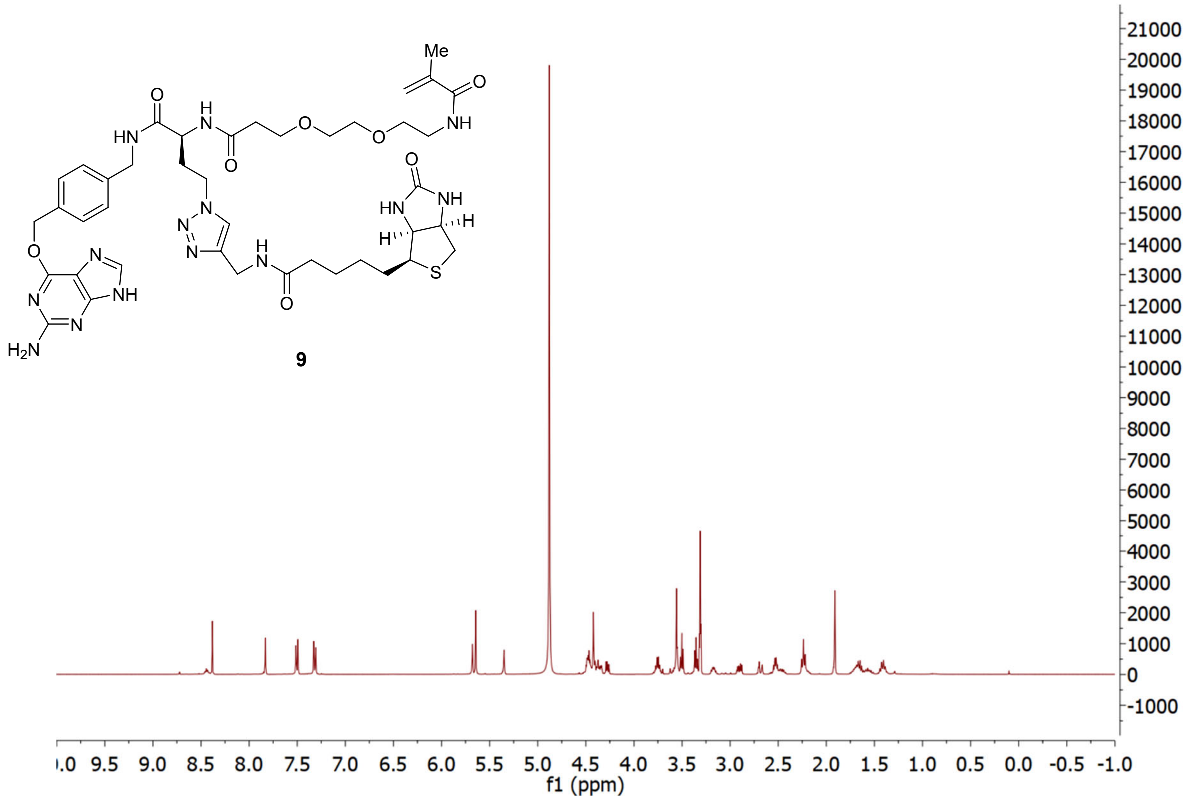


9

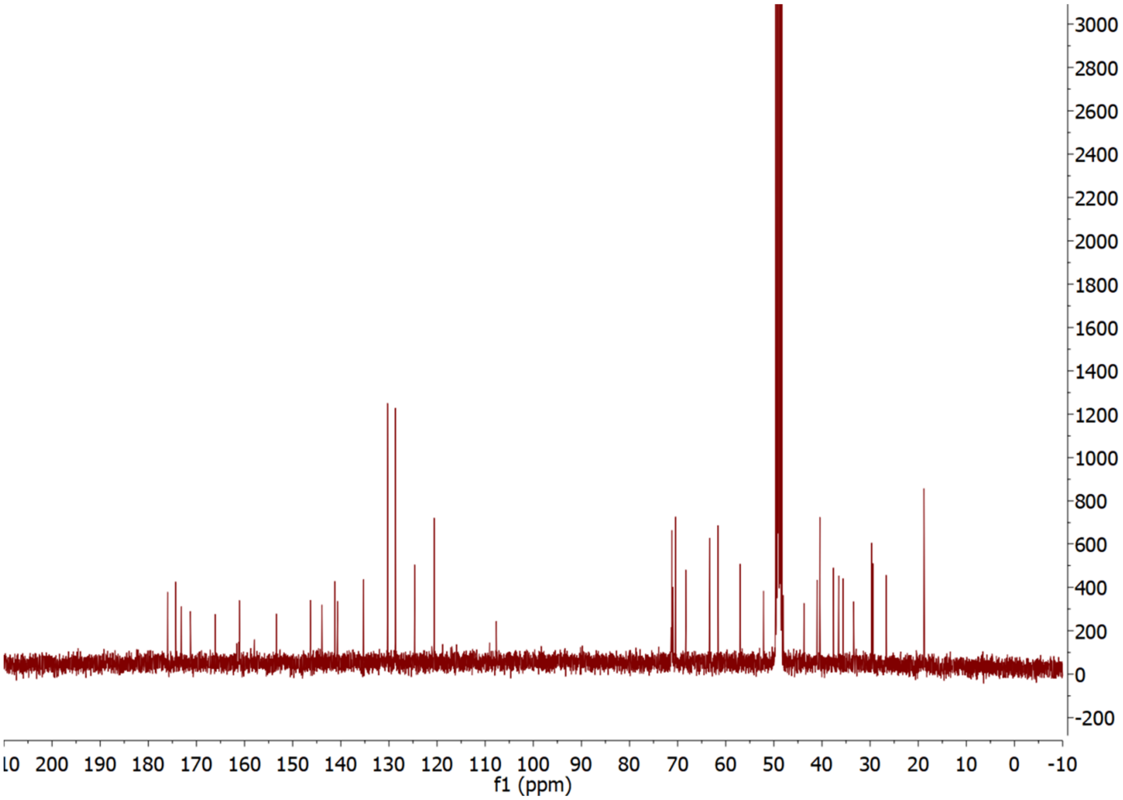


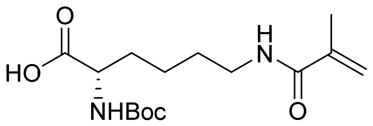


**10**

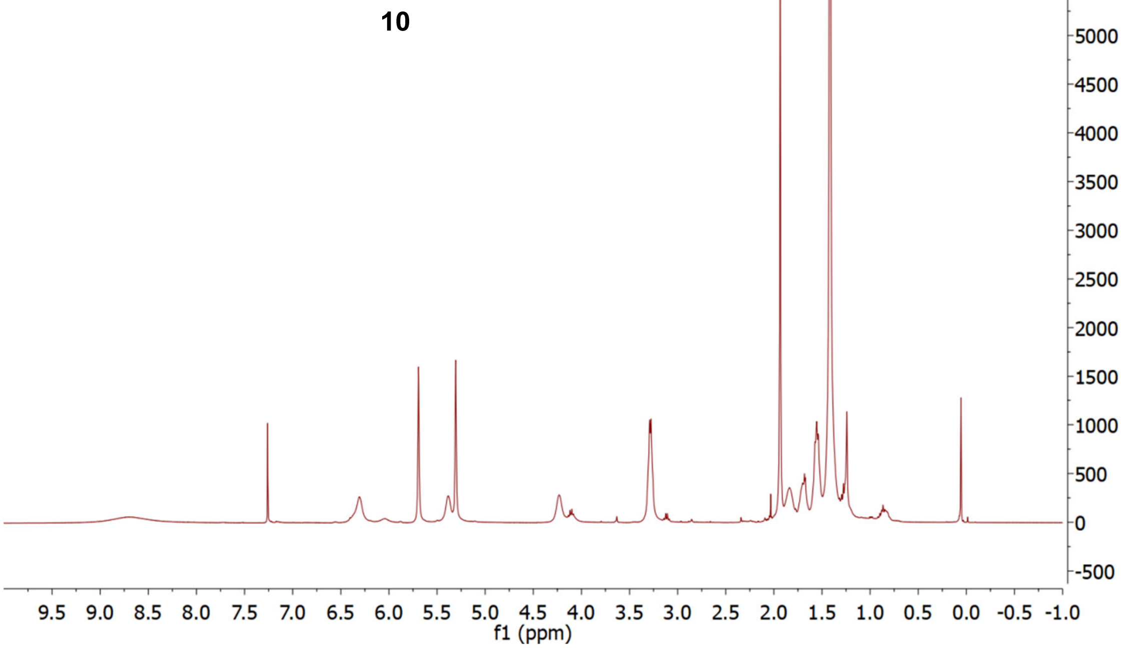

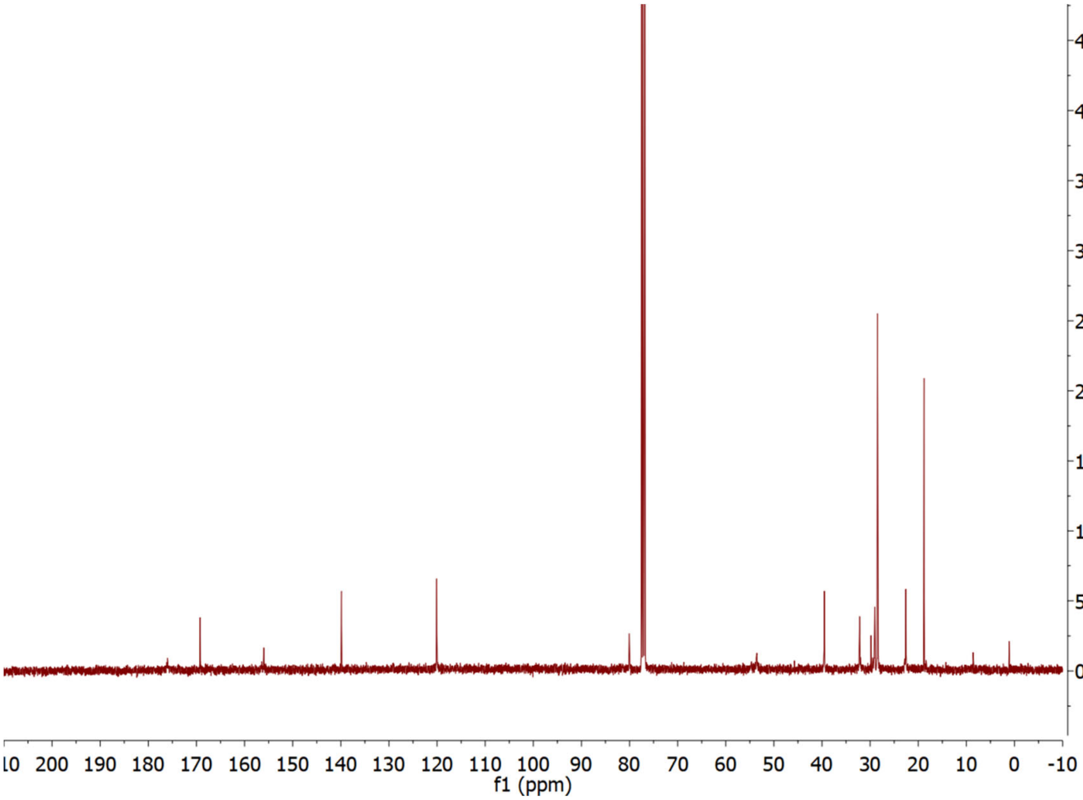
f1 (ppm)

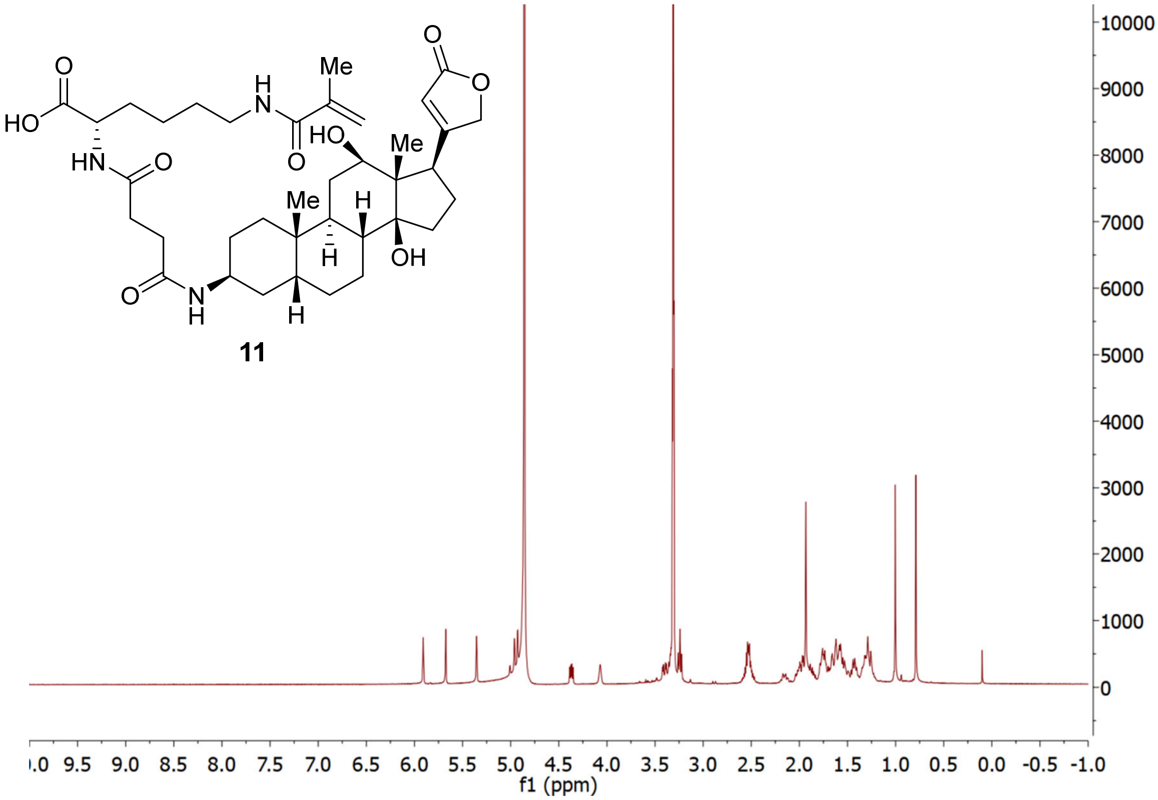
11
f1 (ppm)

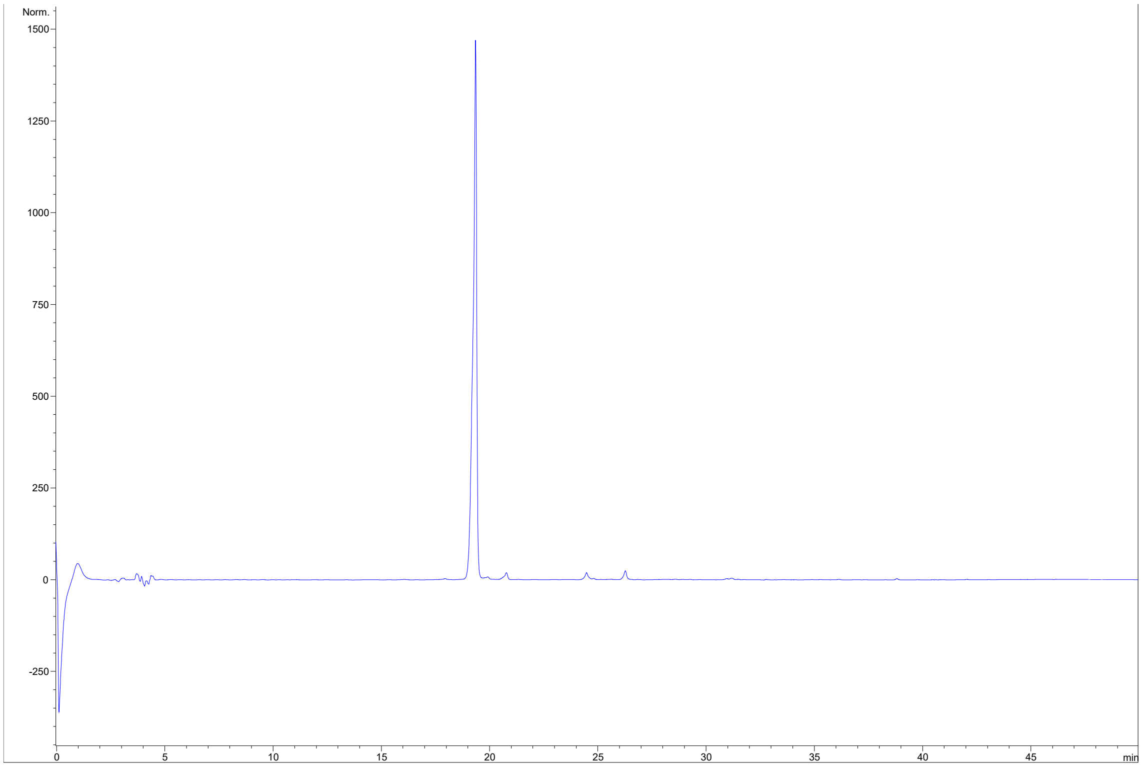


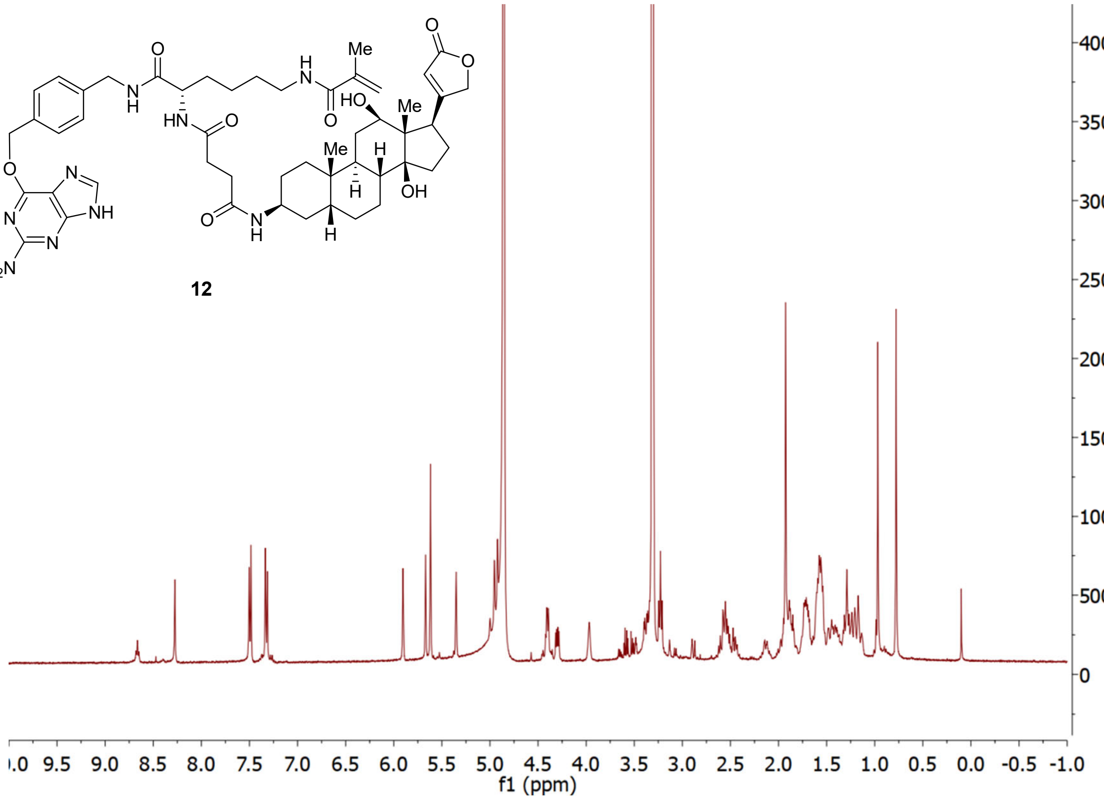


**12**

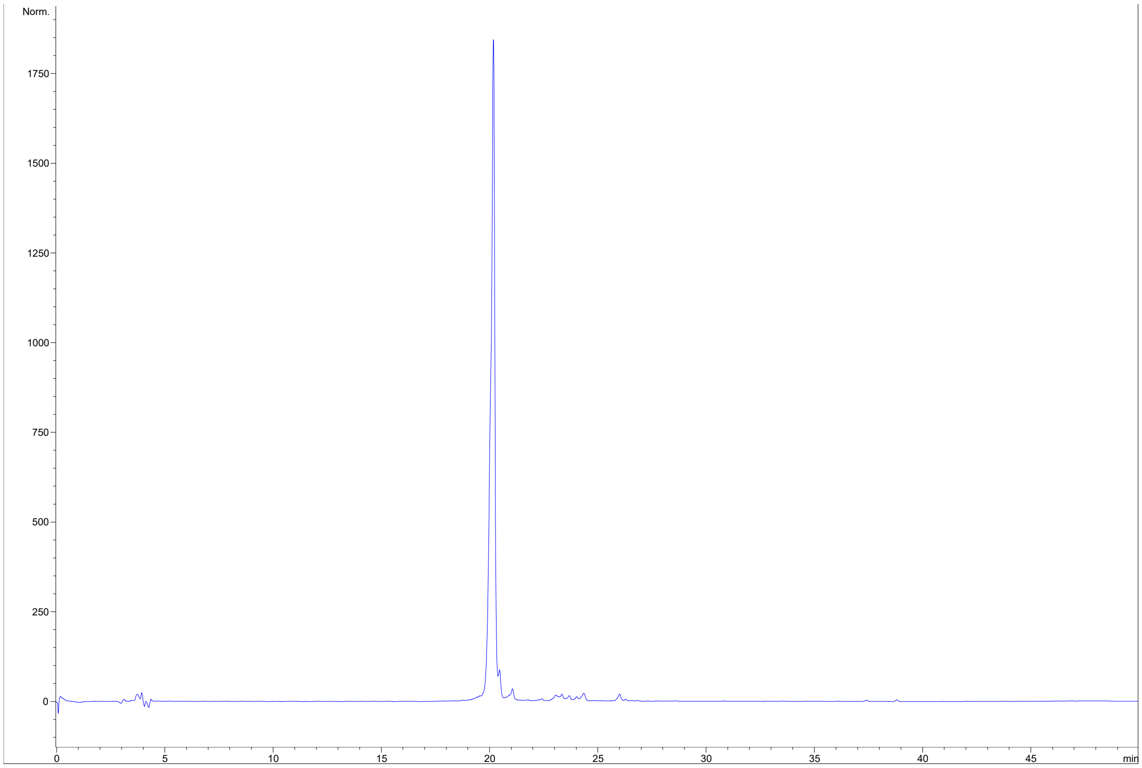
Norm.
1750
1500
1250
1000
750
500
250
0
0
5
10
15
20
25
30
35
40
45
min

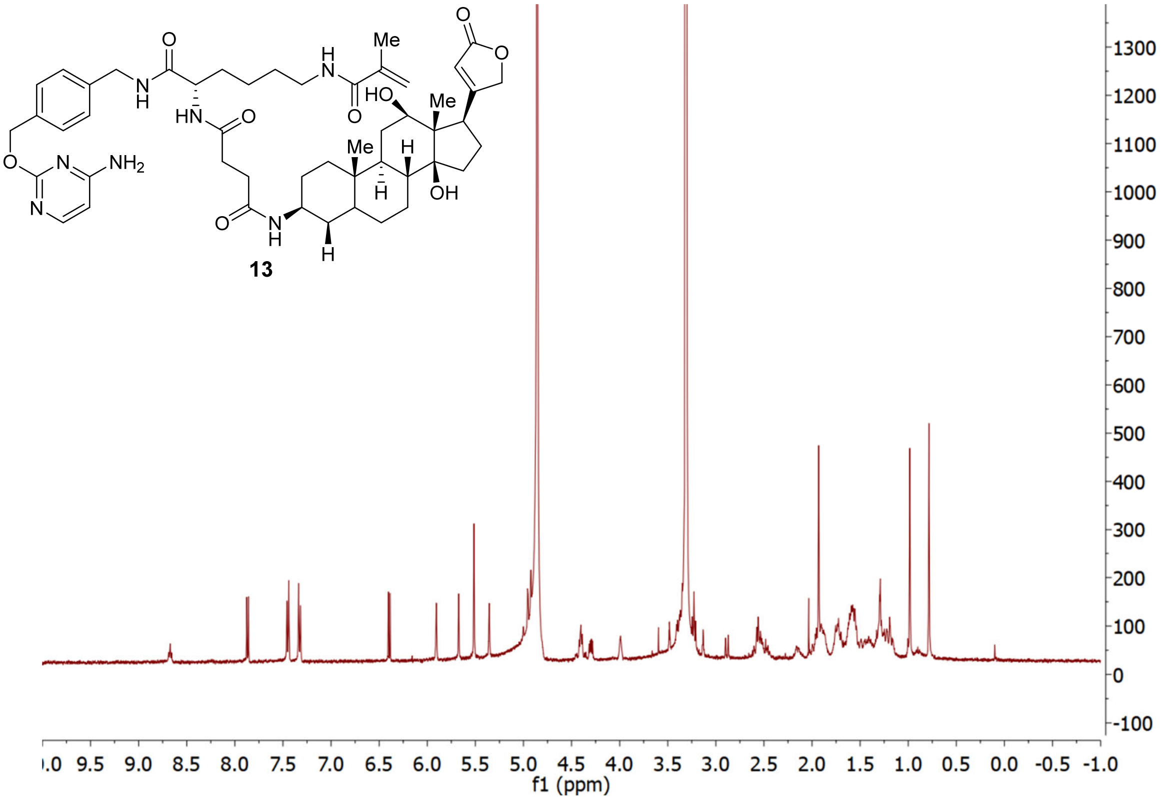
Me
O
O
O
HO
Me
H
Me
H
OH
OH
NH2
N
N
O
H
N
HN
H
N
O
O
O
H
N
H
13
1300
1200
1100
1000
900
800
700
600
500
400
300
200
100
0
-100
f1 (ppm)

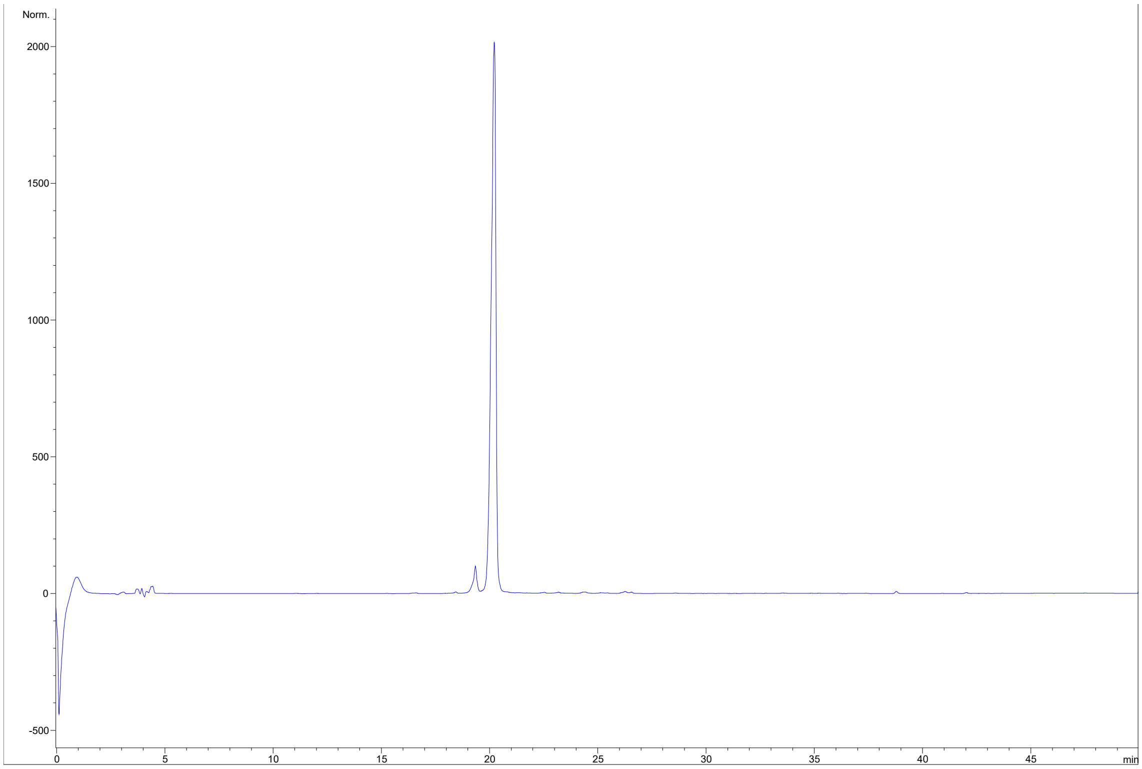

Norm.
2000
1500
1000
500
0
-500
0
5
10
15
20
25
30
35
40
45
min
